# Differentiable Gene Set Enrichment Analysis for Pathway-Level Supervision in Transcriptomic Learning

**DOI:** 10.64898/2026.03.18.712610

**Authors:** Shuaiyu Li, Yang Ruan, Yang Xinyue, Wen Zhang, Hiroto Saigo

## Abstract

In transcriptomics-driven drug discovery, upstream predictors of chemical-induced transcriptional profiles (CTPs) are typically trained with gene-wise objectives, whereas downstream interpretation relies on pathway-level, rank-based statistics such as Gene Set Enrichment Analysis (GSEA). This objective mismatch destabilizes pathway conclusions under prediction errors: small ranking perturbations can flip enrichment direction or distort pathway ordering. To bridge this gap, we present differentiable GSEA (dGSEA), a training-compatible surrogate that maps predicted gene-level scores to pathway enrichment with well-behaved gradients. Technically, dGSEA replaces discrete ranking operations with temperature-controlled soft sorting, smooth prefix accumulation, and differentiable extremum aggregation. Critically, to preserve the statistical semantics of classical GSEA, we introduce sign-specific robust permutation normalization (dNES) with optional *κ*-calibration. For computational efficiency, a scalable Nyström–window approximation (nyswin) reduces the quadratic bottleneck to near-linear complexity, enabling genome-scale evaluation. Empirically, across synthetic benchmarks and LINCS L1000 signatures, dGSEA matches classical GSEA accuracy with improved numerical stability. When incorporated as an auxiliary objective for SMILES-to-transcriptome prediction, dGSEA improves pathway-level agreement (macro correlation 0.257 → 0.306; sign accuracy 0.620 → 0.641) without compromising gene-level performance, providing a practical mechanism for pathway-aware optimization in transcriptomic prediction pipelines.

## 1 Introduction

Predicting chemical-induced transcriptional profiles (CTPs) from molecular structure representations, such as Simplified Molecular Input Line Entry System (SMILES) strings, has emerged as a central task in phenotype-based drug discovery and mechanism-of-action studies [16, 17, 24, 26]. A growing body of deep learning methods has been proposed to model drug-induced gene expression changes directly from small-molecule structures [14, 15, 17, 26, 31, 32, 36], spanning diverse architectures such as graph neural networks [6], Transformer-based encoders [5, 13], and variational autoencoders [7, 11]. Despite this architectural diversity, nearly all existing methods are trained and evaluated using gene-level regression objectives, such as mean squared error (MSE) or Pearson correlation, which implicitly treat all genes as equally important during optimization.

This gene-centric paradigm conflicts with how predicted transcriptional signatures are actually interpreted in downstream applications. Hypotheses in drug repurposing and mechanism-of-action studies are rarely formulated at the level of individual genes; instead, they rely on pathway-level analyses such as gene set enrichment analysis (GSEA) [21] and connectivity scoring [12, 22], which assess coordinated expression patterns across biologically defined gene sets. Critically, because pathway-level statistics such as the normalized enrichment score (NES) in GSEA are fundamentally rank-based, even small but systematic errors in gene ranking can lead to incorrect pathway activation calls or reversed enrichment directions [19, 35]. Recent studies have shown that models achieving high genome-wide *R*^2^ or low MSE may still fail to recover key differentially expressed genes (DEGs) or biologically meaningful signals [1, 34], underscoring that strong gene-level performance does not guarantee pathway-level reliability.

This structural mismatch between training objectives and downstream decision-making (Fig. 1a) would not be critical if transcriptional profiles could be predicted with near-perfect accuracy. However, both classical [36] and state-of-the-art models [32] for *in silico* perturbation prediction remain far from this regime, with genome-wide Pearson correlations typically not exceeding 0.4–0.5 on standard benchmarks. Under such constraints, models with limited representational capacity must implicitly allocate predictive resources across genes. When optimization does not explicitly emphasize biologically relevant gene sets, capacity may be preferentially directed toward genes that dominate global error metrics rather than those driving pathway-level inference. Consequently, improvements in gene-level metrics can mask substantial functional inconsistencies at the pathway or disease-mechanism level. This motivates training paradigms that actively guide model capacity toward pathway-level coherence, rather than treating pathway analysis as a purely post hoc diagnostic.

**Figure 1:**
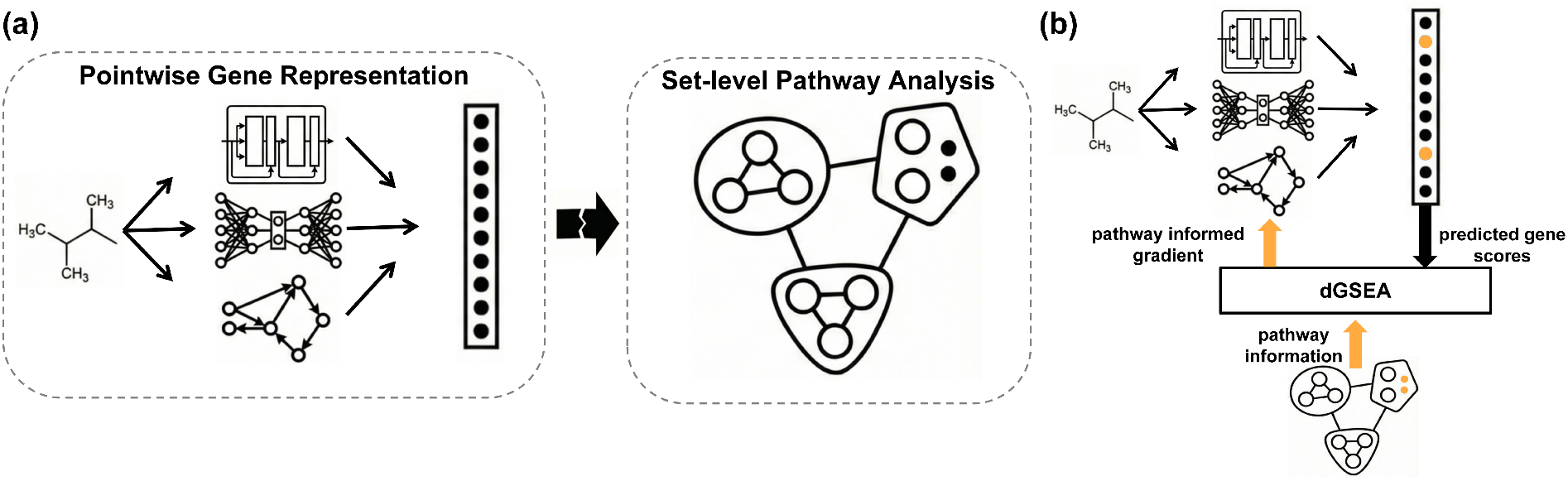
Overview of pathway-aware supervision via dGSEA. **(a)** A mismatch between gene-level training objectives and pathway-level interpretation. **(b)** dGSEA bridges this gap by enabling differentiable, pathway-informed supervision.

To address this mismatch (Fig. 1a), we propose *differentiable GSEA* (**dGSEA**), a smooth surrogate that maps predicted transcriptional profiles to pathway-level enrichment scores with stable gradients, enabling pathway analysis to serve as an explicit training signal rather than a post hoc diagnostic. This approach transforms GSEA from a purely interpretive tool into a training objective directly aligned with the downstream workflows used in drug repurposing and mechanism-of-action studies.

Classical GSEA is incompatible with gradient-based training due to its reliance on non-differentiable operations, including hard ranking and extremum selection. Constructing a differentiable surrogate that can serve as a training objective poses three technical challenges: replacing discrete operations with smooth, gradient-stable relaxations; preserving semantic alignment with the classical NES scale to maintain biological interpretability; and achieving computational efficiency for repeated genome-scale evaluation during training, where classical GSEA’s post hoc complexity becomes prohibitive. dGSEA addresses these challenges through three coordinated mechanisms. Technically, it replaces discrete operations with temperature-controlled soft sorting and smooth prefix kernels. To preserve the statistical semantics of classical GSEA, we introduce sign-specific robust permutation normalization (dNES) with optional *κ*-calibration to the classical null distribution. For computational efficiency, we develop a scalable Nyström-based sliding-window approximation (`nyswin`) that reduces complexity to near-linear, enabling genome-scale end-to-end training.

We validate dGSEA across synthetic benchmarks and the LINCS L1000 dataset [12], where it closely matches classical GSEA while exhibiting improved numerical stability. When incorporated as an auxiliary structured loss for SMILES-to-transcriptome prediction (Fig. 1b), dGSEA preserves gene-level accuracy (mean Pearson correlation 0.449 → 0.452; RMSE 0.420 → 0.418) while substantially improving pathway-level agreement: macro pathway correlation increases from 0.257 to 0.306, sign consistency from 0.620 to 0.641, and pathway MSE decreases from 1.784 to 1.610. These improvements are observed across diverse functional contexts including stress response, proteostasis, and cell-cycle control pathways, demonstrating that pathway-aware optimization enhances biological interpretability without sacrificing predictive fidelity.

## 2 Related Work

Existing work spans two complementary but disconnected strands: upstream models that predict gene-level transcriptional responses, and downstream methods that interpret these predictions through pathway enrichment. Our work bridges this gap by developing a differentiable pathway-level functional that enables end-to-end optimization. We review these two strands to clarify (i) why existing upstream models, despite architectural diversity, share a common limitation in objective design, and (ii) why existing downstream methods, despite statistical rigor, remain incompatible with gradient-based training.

### 2.1 Upstream: Structure-to-Transcriptome Prediction Models

A growing body of work predicts chemical-induced transcriptional profiles (CTPs) from molecular structure, with the LINCS L1000 landmark genes (*G*=978) serving as a common benchmark. Early approaches such as DLEPS [36] established feasibility by encoding SMILES with latent generative models (e.g., GrammarVAE) followed by regression layers. Subsequent methods have strengthened molecular representations through graph neural networks [14, 31, 32] and introduced context conditioning on cell identity, dose, and baseline expression to model condition-specific responses. Notably, DeepCE and MultiDCP incorporate attention mechanisms and multi-task learning to improve cross-cellular generalization [14, 31], while MiTCP injects gene–gene structure from co-expression networks to support coordinated transcriptional prediction [32]. Beyond value regression, several models reformulate the task as differential expression ordering: CIGER and BADGER cast structure-to-signature prediction as a learning-to-rank problem, emphasizing relative gene rankings and introducing pathway- or network-informed inductive biases [9, 15]. More recently, conditional generative models such as TranSiGen and PRnet employ variational frameworks to capture perturbation signals under noise and heterogeneity, scaling to larger perturbation corpora and broader cellular coverage [17, 26].

Despite these architectural advances spanning graph encoders, Transformers, variational autoencoders, and learning-to-rank frameworks all aforementioned methods optimize gene-level objectives, whether regression losses, ranking concordance, or reconstruction likelihoods, without explicit pathway-level constraints. Pathway enrichment, when performed, remains a post hoc interpretive step rather than a training-time objective. Consequently, these models inherit the objective mismatch identified in Section 1: strong gene-level metrics do not guarantee pathway-level fidelity, particularly under the limited prediction accuracy characteristic of current structure-to-transcriptome models.

### 2.2 Downstream: Pathway-level Enrichment and Interpretation Methods

While upstream models predict gene-level outputs, downstream interpretation relies on pathway enrichment methods that aggregate expression signals across gene sets. Classical GSEA provides the canonical framework: genes are ranked by a univariate statistic, and pathway enrichment is quantified via a Kolmogorov–Smirnov-like running-sum score, with significance and normalized enrichment scores assessed through permutation testing [21]. Numerous variants extend this paradigm. Single-sample methods such as ssGSEA and GSVA produce persample pathway scores by comparing in-set and out-of-set gene distributions, enabling pathway-by-sample matrices for downstream modeling [3, 8]. Other approaches replace explicit ranking with continuous aggregations: PLAGE and PAGE summarize gene-set activity using singular-vector decomposition or z-score statistics [10, 25], while CAMERA and ROAST adjust for inter-gene correlation through competitive testing or rotation-based resampling [29, 30].

Despite this methodological diversity, pathway enrichment methods were developed as post hoc inferential tools applied after expression profiles are obtained. They rely on discrete ranking, resampling, or global statistical estimation, which precludes integration into gradient-based optimization. As a result, no existing approach unifies pathway-aware interpretation with differentiable training objectives: upstream predictors optimize pathway-agnostic gene-level losses, while downstream enrichment methods provide biologically meaningful but non-differentiable pathway statistics. dGSEA bridges this gap by constructing a pathway-level functional that preserves the rank-based semantics of classical GSEA through calibrated normalization while enabling end-to-end optimization via smooth relaxations of ranking and aggregation operations, supported by efficient Nyström-based acceleration for genome-scale computation.

## 3 Results

We present results in four stages. First, we formulate differentiable gene set enrichment analysis (dGSEA) as a pathway-level functional suitable for end-to-end optimization. Second, we assess its consistency with classical GSEA and numerical stability. Third, we evaluate scalability and demonstrate training-time feasibility at genome scale. Finally, we apply dGSEA as a structured supervision signal in a SMILES-to-transcriptome prediction task.

### 3.1 Differentiable GSEA as a pathway-level functional

Classical GSEA relies on three non-differentiable operations that preclude integration into gradient-based learning: hard ranking to order genes by scores, discrete prefix accumulation to compute running-sum statistics, and extremum selection to extract maximal enrichment. These operations confine pathway enrichment to post hoc analysis rather than training-time objectives.

We introduce *differentiable GSEA* (dGSEA), a pathway-level functional that maps a gene-level score vector *s* ∈ ℝ^*G*^ and a gene-set indicator *g* ∈ {0, 1}^*G*^ to a scalar enrichment statistic with stable gradients. dGSEA retains the rank-based semantics of classical GSEA by replacing each non-differentiable step with a principled continuous relaxation designed to converge to the classical operation as temperature parameters vanish.

#### Soft ranking

We replace hard ranking with a temperature-controlled soft rank following standard continuous ranking relaxations [18]. Using the sigmoid *σ*(*z*) = (1 + *e*^−*z*^)^−1^ and temperature *τ*_rank_ *>* 0, the soft rank of gene *i* is defined as

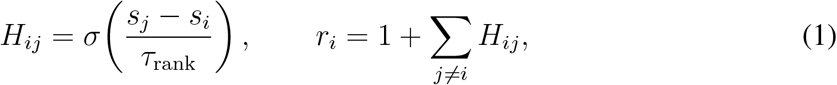

where *r*_*i*_ continuously approximates the descending rank of gene *i*. As *τ*_rank_ → 0, *H*_*ij*_ approaches a step function, recovering hard ranking.

#### Smooth prefix accumulation

To construct a differentiable running-sum curve, we introduce a soft prefix indicator with temperature *τ*_prefix_ *>* 0. For each position *t* ∈ {1, …, *G*} and gene *i*,

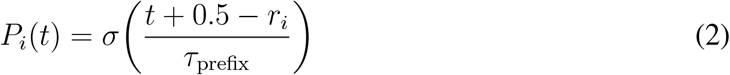

softly selects whether gene *i* lies within the top-*t* positions. Accumulating weighted hit and miss contributions yields

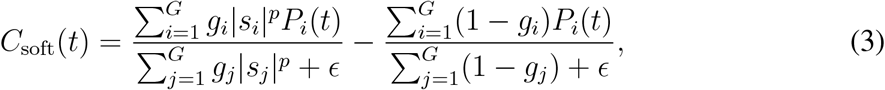

where *p* ≥ 0 is the weighting exponent (typically *p* = 1) and *ϵ* is a numerical stability constant. This provides a smooth surrogate of the classical running-sum statistic.

#### Differentiable extremum aggregation

Classical GSEA extracts the maximum absolute deviation of the running sum via max_*t*_ |*C*_*t*_| or min_*t*_ *C*_*t*_, which is non-differentiable at ties or inflection points. We replace this with a temperature-weighted aggregation. With *τ*_abs_ *>* 0,

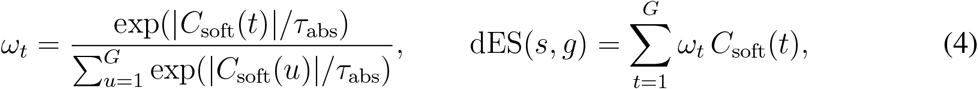

where *ω*_*t*_ assigns higher weight to positions with larger absolute deviations. As *τ*_abs_ → 0, this converges to selecting the position with maximal |*C*_soft_(*t*)|.

Fig. 2 provides a schematic overview of the dGSEA pipeline. The resulting differentiable enrichment score dES(*s, g*) serves as a smooth surrogate of the classical enrichment score, enabling pathway enrichment to be evaluated within gradient-based learning pipelines.

**Figure 2:**
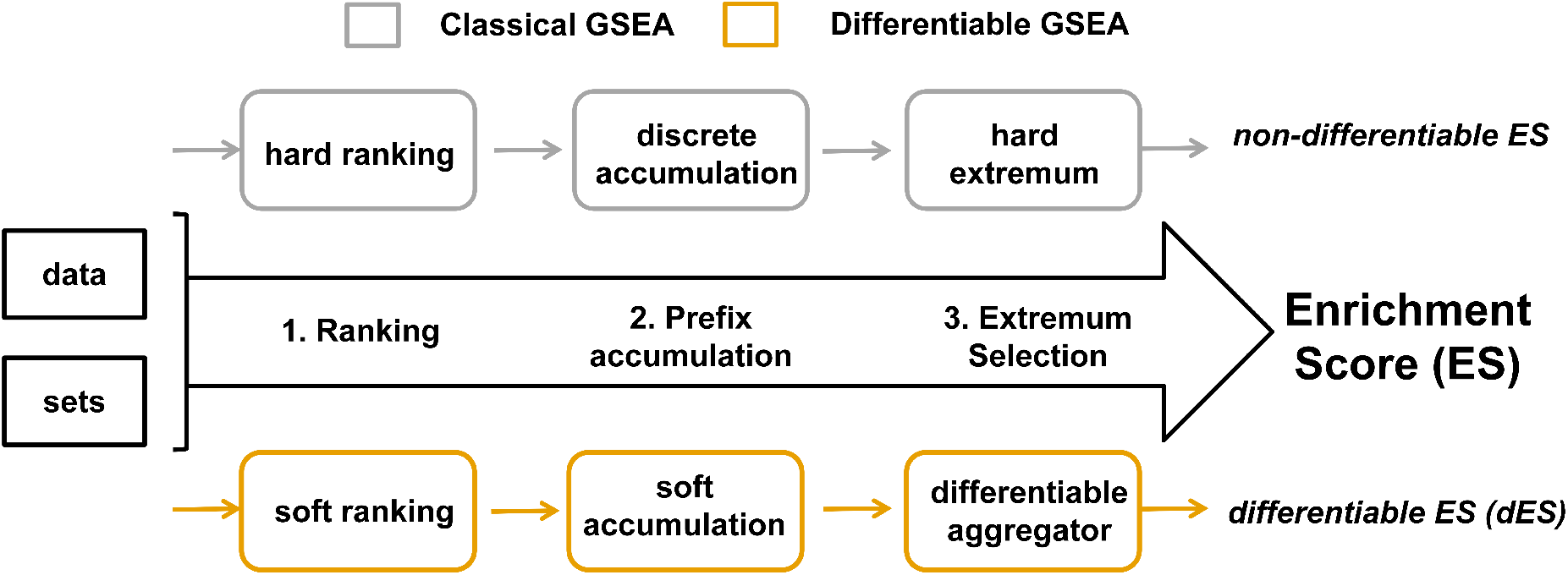
Schematic overview of dGSEA. Classical GSEA operations (hard ranking, discrete prefix accumulation, and extremum selection) are replaced by smooth, differentiable counterparts. The resulting pathway-level functional produces a differentiable enrichment score (dES), enabling enrichment to be evaluated and optimized within gradient-based learning pipelines.

#### Theoretical guarantees

These relaxations are not heuristic approximations. As the temperatures vanish *τ*_rank_, *τ*_prefix_, *τ*_abs_ → 0 the differentiable enrichment score converges pointwise to the classical enrichment score under the assumption of distinct gene scores and a unique extremum (Theorem B.1), establishing that dGSEA is a consistent estimator of classical GSEA in the zero-temperature limit. Moreover, dGSEA is continuous everywhere and differentiable almost everywhere in *s*, with gradients admitting explicit bounds that scale with the temperature parameters (Proposition B.6). These bounds decompose the gradient into a *direct* effect through the weighted hit term and a *rank-mediated* effect induced by soft ranking and prefix kernels, providing a principled explanation for why dGSEA yields controlled, training-compatible gradients.

Fig. 3 compares classical and differentiable running-sum curves across eight controlled synthetic scenarios spanning mild responses, heavy-tailed noise, targeted perturbations, broad-spectrum effects, batch shifts, and dose-dependent responses. Even at finite temperatures and under realistic noise conditions, the smooth accumulation preserves the characteristic trajectory and enrichment direction of classical GSEA while avoiding discontinuities induced by hard ranking and extremum selection. Detailed scenario definitions are provided in Appendix A.1. Unless otherwise specified, we use the default temperature setting

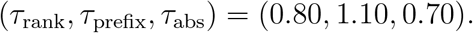

**Figure 3:**
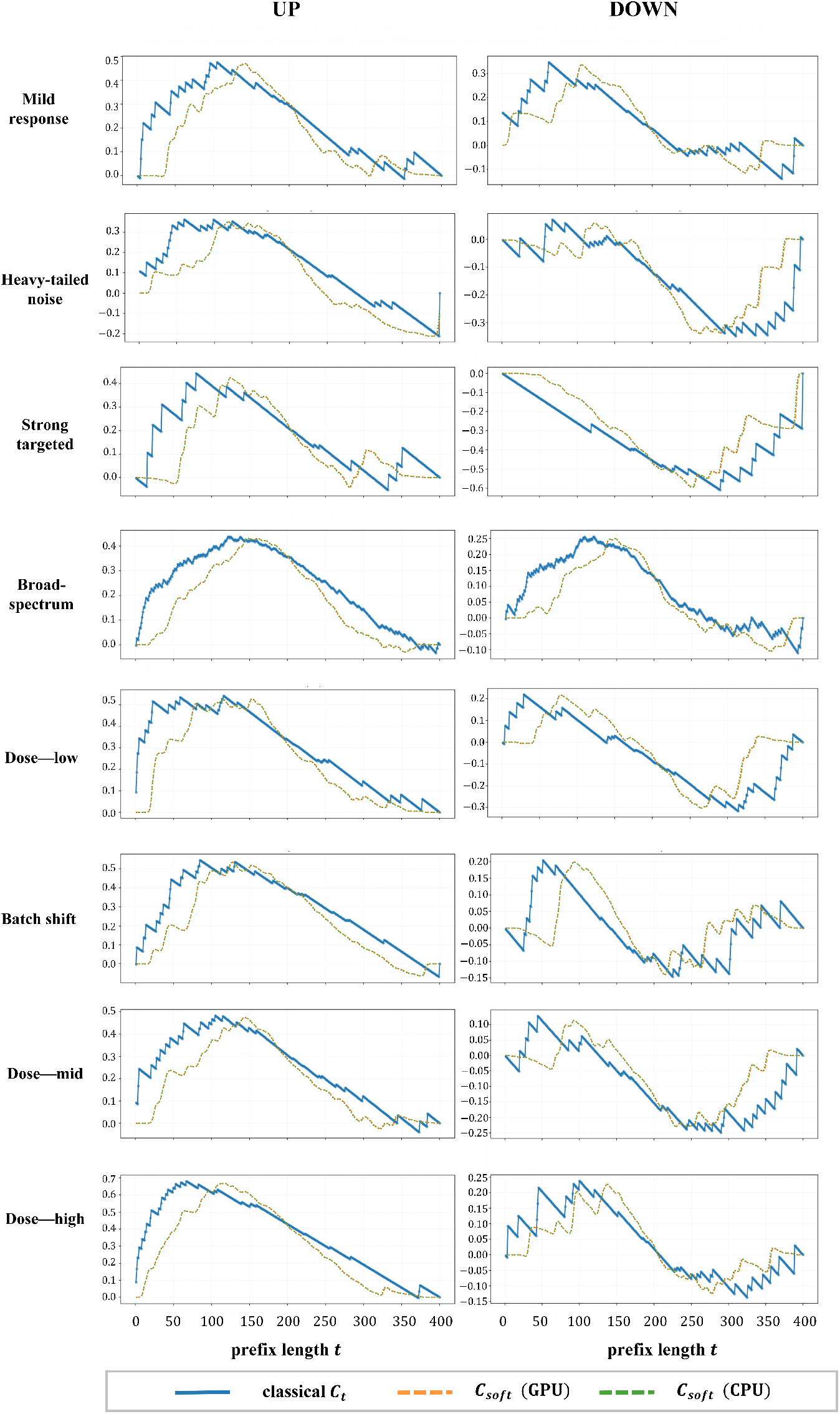
Representative running-sum curves comparing classical GSEA (*C*_*t*_, discrete step function) and dGSEA (*C*_soft_(*t*), smooth curve) across eight controlled synthetic scenarios with varying signal strength, gene-set size, and noise characteristics. Each panel shows paired UP (enrichment, left) and DOWN (depletion, right) gene sets. Scenarios span mild responses (top), heavy-tailed noise, targeted perturbations, broad-spectrum effects, batch shifts, and dose-dependent responses (bottom). The smooth curves preserve the characteristic trajectory and enrichment direction of classical GSEA while avoiding discontinuities. Detailed scenario definitions are provided in Appendix A.1.

Sensitivity analyses and recommended parameter ranges are provided in Appendix C.

Since practical interpretation of pathway activity relies on the normalized enrichment score (NES), we next examine whether dGSEA recovers the statistical semantics of NES and whether this alignment improves numerical stability.

### 3.2 Recovering classical GSEA semantics with improved stability

A differentiable surrogate is practically useful only if it preserves the statistical semantics and qualitative behavior that make classical GSEA interpretable. We evaluate dGSEA with respect to three core properties that collectively establish its validity as a surrogate for classical GSEA: *statistical fidelity* (recovery of classical enrichment scores and valid permutation *p*-values under the null), *scale alignment* (preservation of sign and relative ordering compared to classical NES), and *numerical stability* (reduced sensitivity to input perturbations compared to hard ranking and extremum selection). We validate these properties using controlled synthetic benchmarks and real transcriptomic data from the LINCS L1000 dataset. Detailed experimental settings are provided in Appendices A.1 and A.2.

#### Directional pathways and sign-specific normalization

Many biological pathways exhibit *directional regulation*, where distinct subsets of genes are coordinately up-regulated or down-regulated in response to perturbations. We therefore evaluate dGSEA on *directional pathways* defined by paired gene-set indicators: an UP set *g*_up_ ∈ {0, 1}^*G*^ and a DOWN set *g*_dn_ ∈ {0, 1}^*G*^. Given a score vector *s* ∈ ℝ^*G*^, classical GSEA computes enrichment scores ES_up_ and ES_dn_ for each component and normalizes each by the expected magnitude of its permutation null distribution. Because the weighted running-sum formulation assigns different effective weights to genes inside versus outside the set, the resulting permutation null distribution can be *sign-asymmetric*, particularly when gene-set size deviates substantially from *G/*2:

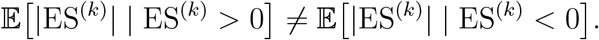

To account for this asymmetry, classical GSEA employs sign-specific normalization: NES = ES*/µ*_*σ*_, where *σ* = sign(ES) and *µ*_*σ*_ is the mean absolute enrichment score over permutations with sign *σ*.

dGSEA follows this normalization principle in a fully differentiable manner. For each directional component, we compute a smooth enrichment score dES and estimate the sign-conditional null scale from *K* = 400 gene-label-permuted score vectors. To limit sensitivity to extreme permutation values, we use a robust mean estimator that combines a symmetric *ρ*-trimmed mean and a *ρ*-Winsorized mean. Here, *ρ* ∈ (0, 1*/*2) denotes the trimming proportion controlling the fraction of extreme permutation values removed from each tail of the null distribution, and *λ* ∈ [0, 1] is a convex mixing weight that interpolates between the trimmed and Winsorized estimators. Unless otherwise specified, we fix *ρ* = 0.15 and *λ* = 0.10 in all experiments:

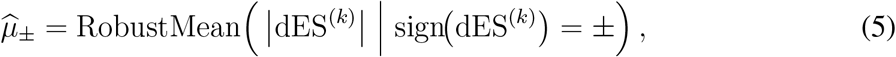

where 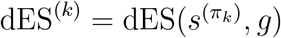 denotes the enrichment score for the *k*-th permutation *π*_*k*_ (Proposition B.4). The normalized statistic is then 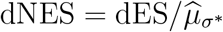, where *σ*^∗^ = sign(dES).

When benchmarking dGSEA against classical GSEA (Table 1), we additionally apply *κ*- calibration (Proposition B.5) to align the dNES scale with classical NES, facilitating direct numerical comparison. This calibration is omitted when using dGSEA as a training objective, where relative pathway ordering rather than absolute scale is critical.

**Table 1:** Comparison of classical GSEA and dGSEA across eight synthetic scenarios. Normalized enrichment scores (NES, dNES) are shown for UP and DOWN gene sets, along with scale compression ratios (dNES/NES) and Spearman correlation between running-sum curves. dNES values are computed without *κ*-calibration.

| Scenario | Normalized Enrichment Score |  |  |  | Scaling |  | Curve |  |
| --- | --- | --- | --- | --- | --- | --- | --- | --- |
|  | Classical |  | dGSEA |  | dNES/NES |  | Spearman |  |
|  | UP | DN | UP | DN | UP | DN | UP | DN |
| Mild response | 1.64* | 1.25 | 0.67* | 0.31 | 0.41 | 0.25 | 0.73 | 0.74 |
| Strong targeted | 1.30 | -1.77** | 0.40 | -0.91** | 0.31 | 0.52 | 0.57 | 0.85 |
| Broad-spectrum | 1.91** | 1.28* | 0.77** | 0.23* | 0.41 | 0.18 | 0.83 | 0.79 |
| Heavy-tailed | 1.23 | -1.13 | 0.16 | -0.39 | 0.13 | 0.35 | 0.89 | 0.58 |
| Batch shift | 1.62** | 0.80 | 0.79** | 0.05 | 0.48 | 0.06 | 0.81 | 0.28 |
| Dose (low) | 1.85** | -1.11 | 0.94** | -0.10 | 0.51 | 0.09 | 0.72 | 0.56 |
| Dose (mid) | 1.78* | -0.92 | 0.72** | -0.20 | 0.40 | 0.21 | 0.78 | 0.66 |
| Dose (high) | 2.39** | 0.84 | 1.26** | 0.11 | 0.53 | 0.13 | 0.76 | 0.72 |

Significance levels: \* $p < 0.05$ ; \*\* $p < 0.01$ , from one-sided permutation tests ( $K=400$ gene-label permutations) assessing enrichment under the null of no gene-set association. P-values are uncorrected, as this table serves a diagnostic purpose comparing dGSEA and classical GSEA behavior across synthetic regimes. The ratio dNES/NES quantifies scale compression.

#### Synthetic benchmarks

We first validate these properties under fully controlled settings where ground-truth enrichment structure is known. Score vectors are simulated over a fixed gene universe (*G* = 400) with paired UP/DOWN gene sets evaluated across eight regimes spanning weak to strong signals, narrow to broad gene-set support, Gaussian and heavy-tailed noise, and batch-like location/scale shifts (full parameterization in Appendix A.1). For each scenario, we generate *M* = 32 independent instances. Classical GSEA and dGSEA are evaluated using identical permutation indices to ensure strictly paired comparisons.

Fig. 4 summarizes the validation. Panel (a) demonstrates scale alignment: dNES exhibits strong linear agreement with classical NES, with an ordinary least squares (OLS) slope of *y* = 0.91*x*. This mild scale compression reflects the distributional nature of soft aggregations soft ranking and softmax-weighted accumulation distribute probability mass across multiple positions, reducing peak magnitude compared to hard extremum selection while preserving rank concordance (Spearman *ρ* = 0.87). Scenario-level results in Table 1 show that dNES/NES ratios typically fall in the range 0.3–0.5 before *κ*-calibration, with consistent preservation of enrichment direction and statistical significance. Importantly, running-sum trajectories remain largely aligned (median curve correlation *ρ* = 0.74, range 0.28–0.89), confirming that relative pathway rankings the primary basis for biological interpretation are maintained even when absolute magnitudes differ.

**Figure 4:**
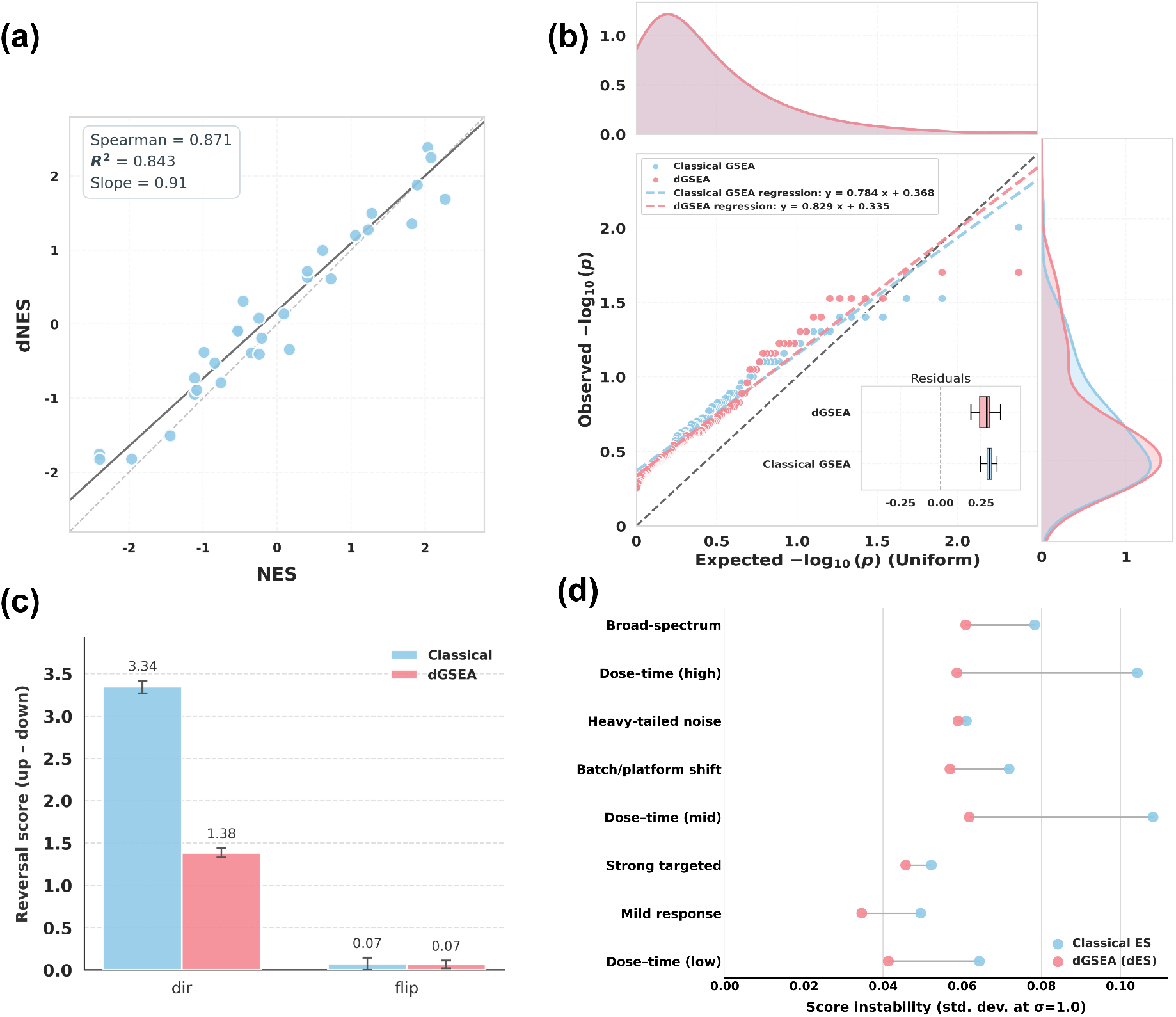
Synthetic benchmarks: semantic agreement, validity and robustness of dGSEA. (a) dNES agrees closely with classical NES with mild scale shrinkage (OLS *y* = 0.91*x*). (b) Under the null, permutation *p*-values are well calibrated in QQ space, with dGSEA slightly closer to uniform and reduced residual dispersion. (c) Two-set reversal preserves opposing-signal interpretability: direct RS is strongly positive, while flip RS collapses to ≈ 0 (dGSEA more conservative in magnitude). (d) Additive-noise stress tests show consistently lower instability for dGSEA across scenarios.

Panel (b) validates statistical fidelity: under the global null, permutation *p*-values closely follow the uniform reference in quantile-quantile space, with dGSEA exhibiting reduced residual dispersion compared to classical GSEA. Panel (c) evaluates directional interpretability through a two-set reversal test, where direct contrasts yield strongly positive scores while flipped-sign controls collapse near zero, demonstrating correct handling of antagonistic regulation. Panel (d) assesses numerical stability via additive-noise stress tests, where dGSEA shows substantially reduced output variability across regimes (mean instability 0.04 vs. 0.06, representing a 33% reduction).

Together, these results demonstrate that dGSEA faithfully preserves the statistical semantics of classical GSEA including enrichment directions, pathway rankings, and permutation validity while providing improved numerical stability through smooth relaxations and robust normalization.

#### Real-world validation on LINCS L1000

We assess whether the semantic agreement observed under controlled settings persists on real transcriptomic data. Using the LINCS L1000 Level-5 compendium (978 landmark genes), we construct compound-level expression signatures by aggregating profiles sharing the same canonical SMILES across experimental conditions (*N* = 10,555 compounds). We evaluate five curated directional pathways: p53 tumor suppressor, Cell cycle (CDK/Cyclin), Heat-shock (DNAJ/HSP40), Ubiquitin-proteasome (DUBs), and DNA damage (GADD45), each defined by paired UP and DOWN gene sets where applicable. To summarize directional regulation, we define a reversal score

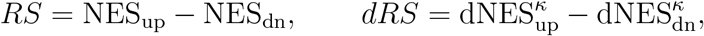

where dNES^*κ*^ denotes *κ*-calibrated dNES aligned to the classical NES scale (Proposition B.5), and *RS* = NES_up_ for pathways lacking a DOWN component.

Across pathways, dGSEA shows strong agreement with classical GSEA in rank-based metrics: Spearman correlations range from *ρ* = 0.91 to 0.98 (median 0.95), and Kendall’s *τ* from 0.75 to 0.86 (Fig. 5a). Absolute rank differences are concentrated near zero (Fig. 5b), and Top-*K* Jaccard overlap increases rapidly with *K*, plateauing at approximately 0.7–0.8 for *K* ≥ 100 (Fig. 5c). While global rankings align closely, Top-50 comparisons reveal a shared core of high-confidence compounds alongside method-specific subsets (Fig. 5d), motivating a deeper analysis of whether these method-specific hits exhibit systematic chemical or functional patterns.

**Figure 5:**
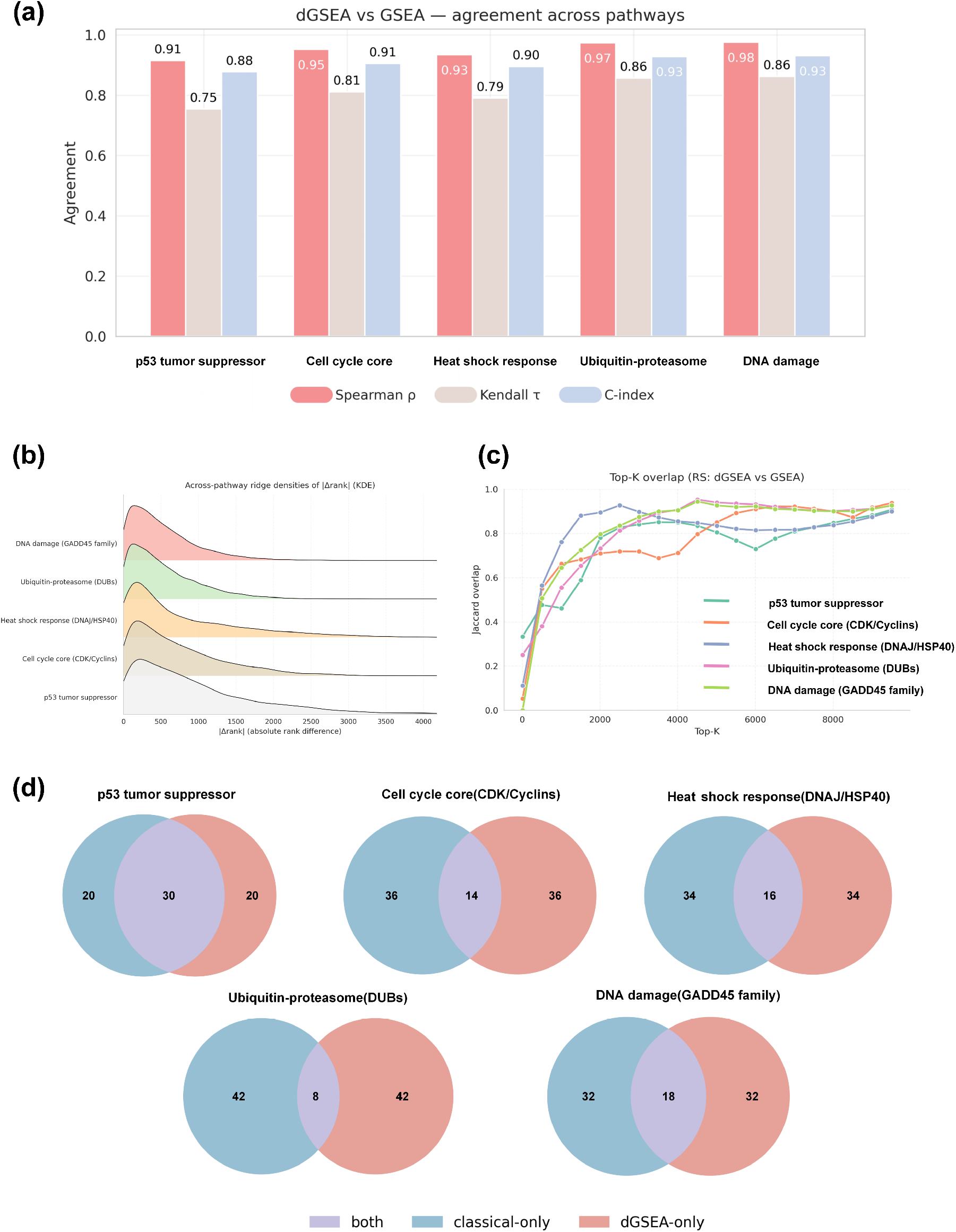
Real-data validation on LINCS L1000: pathway-level agreement and structure of top hits. (a) Across five pathways, directional scores (*RS* vs. *dRS*) show high concordance (Spearman *ρ* ≈ 0.91–0.98; Kendall *τ* ≈ 0.75–0.86). (b) Absolute rank differences are concentrated near zero, indicating limited re-ordering. (c) Top-*K* Jaccard overlap increases with *K* and plateaus at high similarity. (d) Top-50 comparisons reveal a shared core with method-specific tails.

#### Structural coherence of method-specific hits

To assess whether method-specific top-ranked compounds exhibit non-random chemical structure patterns, we partition the Top-50 union for each pathway into three groups *both* (shared by both methods), *classical-only*, and *dGSEA-only* and quantify intra-group chemical homogeneity using mean pairwise Jaccard similarity of Morgan fingerprints (radius=2, 2048 bits). Statistical significance is evaluated via permutation tests (10,000 resamples) with Benjamini-Hochberg FDR correction across 15 comparisons (Table 2).

**Table 2:** Chemical-space homogeneity of Top-50 compound subsets across five pathways. Compounds are ranked by directional enrichment score (RS, dRS) and partitioned into three groups: *both* (shared by both methods), *classical-only*, and *dGSEA-only*. Homogeneity is quantified by mean pairwise Jaccard similarity of Morgan fingerprints (radius=2, 2048 bits). P-values from one-sided permutation tests (10,000 resamples) with Benjamini-Hochberg FDR correction (*α* = 0.05) across 15 comparisons. Bold values indicate significant enrichment after BH FDR correction (adjusted *p <* 0.05).

| Pathway | Group | Count | Observed Jaccard | Null Mean | SD | p-value |
| --- | --- | --- | --- | --- | --- | --- |
| p53 tumor suppressor | both | 30 | 0.1499 | 0.1473 | 0.0066 | 0.338 |
|  | classical-only | 20 | 0.1684 | 0.1474 | 0.0090 | **0.013** |
|  | dGSEA-only | 20 | 0.1396 | 0.1474 | 0.0090 | 0.802 |
| Cell cycle (CDK/Cyclin) | both | 14 | 0.2238 | 0.1614 | 0.0160 | **0.002** |
|  | classical-only | 36 | 0.1706 | 0.1610 | 0.0078 | 0.110 |
|  | dGSEA-only | 36 | 0.1466 | 0.1610 | 0.0078 | 0.975 |
| Heat-shock (DNAJ/HSP40) | both | 16 | 0.2626 | 0.2271 | 0.0270 | 0.103 |
|  | classical-only | 34 | 0.2739 | 0.2264 | 0.0156 | **0.005** |
|  | dGSEA-only | 34 | 0.1916 | 0.2264 | 0.0156 | 0.994 |
| Ubiquitin-proteasome (DUBs) | both | 8 | 0.2159 | 0.2160 | 0.0348 | 0.439 |
|  | classical-only | 42 | 0.1988 | 0.2171 | 0.0112 | 0.961 |
|  | dGSEA-only | 42 | 0.2355 | 0.2171 | 0.0112 | 0.063 |
| DNA damage (GADD45) | both | 18 | 0.1295 | 0.1434 | 0.0117 | 0.902 |
|  | classical-only | 32 | 0.1385 | 0.1440 | 0.0072 | 0.779 |
|  | dGSEA-only | 32 | 0.1655 | 0.1440 | 0.0072 | **0.003** |

Two contrasting patterns emerge. The *classical-only* subsets exhibit significant chemical homogeneity in the p53 (*p* = 0.013) and Heat-shock (*p* = 0.005) pathways, revealing statistical association between classical GSEA’s top rankings and structurally compact chemical clusters in these contexts. Conversely, the *dGSEA-only* subset shows significant homogeneity in the DNA-damage pathway (*p* = 0.003), displaying a similar structural coherence pattern. The *both* (shared) group shows strongest homogeneity in the Cell-cycle pathway (*p* = 0.002), consistent with both methods recovering a common core of well-characterized CDK inhibitors, whereas the Ubiquitin-proteasome pathway shows no significant groupwise differences, reflecting higher mechanistic diversity. Critically, these structural associations emerge even after *κ*-calibration aligns the overall enrichment scale, indicating that they arise from algorithmic differences soft ranking distributes weights across neighboring gene positions, whereas classical GSEA uses hard ranking; similarly, softmax aggregation weights multiple curve positions, whereas classical GSEA selects a single extremum leading to distinct but complementary selection profiles at ranking boundaries.

To contextualize these quantitative findings, we examine functional annotations of representative method-specific compounds visualized in chemical space (Fig. 6). Consistent with the marker legend in Fig. 6, the annotated examples shown in the three panels correspond to *dGSEA-only* selections (red triangles). In the Cell-cycle pathway, the dGSEA-only subset includes known CDK inhibitors such as *Purvalanol A* [27]. In the Heat-shock pathway, the dGSEA-only subset includes stress-response modulators such as *Bortezomib* [20] and *KNK437* [33], as well as *Tipifarnib*. In the DNA-damage pathway, the dGSEA-only subset includes compounds reported to induce GADD45-related responses, including *Anisomycin* [23], *Piperlongumine* [2], and *YM155* [4]. These examples indicate that method-specific selections, while differing from classical GSEA’s top rankings, remain biologically plausible and functionally coherent within their respective pathways; the embeddings are intended as qualitative illustrations consistent with the quantitative homogeneity analysis (Table 2) rather than standalone statistical validation.

**Figure 6:**
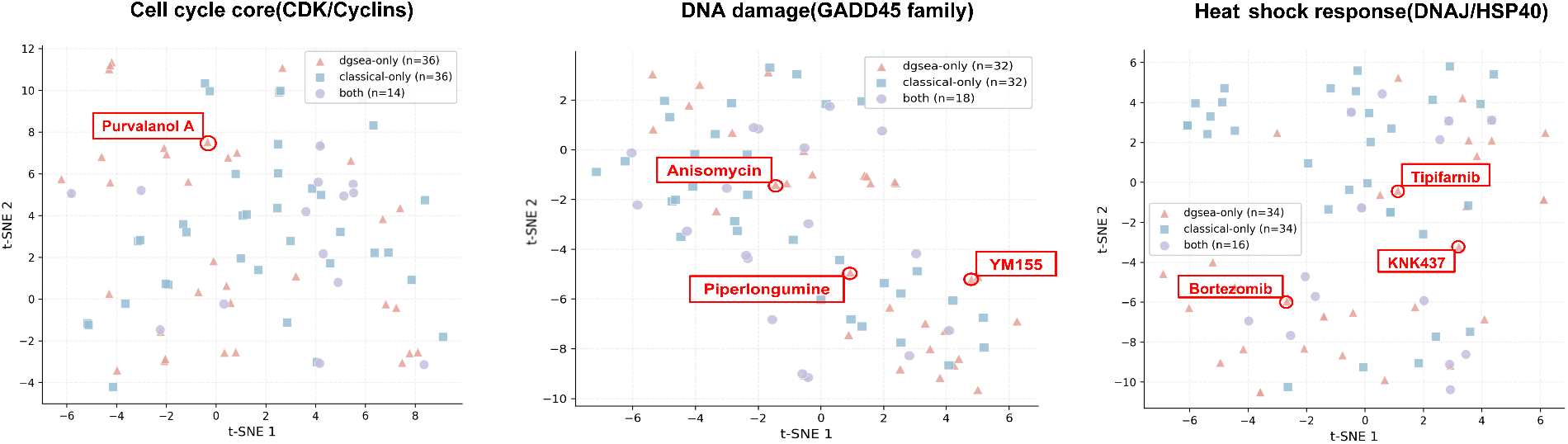
Functional coherence of method-specific hits. t-SNE visualization of Top-50 union compounds across representative pathways illustrates non-random chemical structure in method-specific subsets. Annotated compounds demonstrate functional coherence: Cell-cycle pathway includes CDK inhibitors (e.g., Purvalanol A); Heat-shock pathway includes stress-response modulators (e.g., Bortezomib, KNK437); DNA-damage pathway includes GADD45-related compounds (e.g., Anisomycin, Piperlongumine, YM155). These embeddings provide qualitative illustration consistent with quantitative homogeneity analysis (Table 2).

### 3.3 Scalable implementation enables training-time usage

Having validated dGSEA’s semantic fidelity and numerical stability, we now address computational efficiency. Integrating pathway enrichment into model training requires repeated evaluation at genome scale under tight computational budgets. The naive dGSEA formulation, however, exhibits two quadratic bottlenecks with respect to the number of genes *G*: soft ranking requires all-pairs comparisons (*O*(*G*^2^) time and memory), and smooth prefix accumulation evaluates the running-sum curve across the full rank grid (an additional *O*(*G*^2^) term). While acceptable for small synthetic settings (*G* ∼ 400), this becomes prohibitive for typical transcriptomic profiles (*G* ∼ 10^3^–10^4^) and is incompatible with repeated evaluation inside training loops.

#### Nyström-windowing approximation

To overcome these bottlenecks, we develop `nyswin`, a scalable variant combining Nyström-based anchor sampling for soft ranking with adaptive windowing for prefix accumulation. These two approximations are complementary: Nyström reduces the cost of pairwise comparisons, while windowing restricts evaluation to the rank region expected to contain the extremum.

For soft ranking, the exact formulation computes

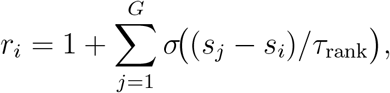

where *σ*(·) is the sigmoid and *τ*_rank_ is the temperature. We approximate this by sampling *m* = 512 ≪ *G* anchors 𝒜_*m*_ = {*a*_1_, …, *a*_*m*_} as uniform quantiles of the score distribution, yielding

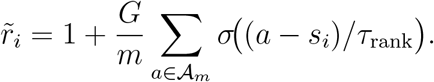

This reduces complexity from *O*(*G*^2^) to *O*(*Gm*), with error bounded by the Kolmogorov– Smirnov discrepancy between the anchor distribution and the full score distribution (Theorem B.7).

For prefix accumulation, we restrict evaluation to a windowed rank grid 𝒯 ⊂ {1, …, *G*} centered around the median soft rank with adaptive half-width *T* ∈ [⌊0.1*G*⌋, *G/*2] determined by gene-set size and rank dispersion, yielding window length |𝒯 | ∈ [0.2*G, G*]. The differentiable enrichment score is computed using softmax-normalized contributions restricted to 𝒯, reducing prefix evaluation cost while preserving the dominant contribution from the extremal region. The approximation error is bounded by the softmax mass outside the window and becomes exact when the window contains all maximizers of the running-sum curve (Proposition B.8).

Together, these approximations reduce dominant quadratic costs to near-linear complexity in *G* while preserving semantic behavior.

#### Approximation fidelity and runtime performance

Table 3 and Fig. 7 validate approximation quality across synthetic scenarios. The accelerated `nyswin` implementation closely matches exact dGSEA, with mean relative error of 0.134% (maximum 0.166%), substantially smaller than the intrinsic variability introduced by permutation-based normalization. Bland– Altman analysis confirms tight distributional agreement, demonstrating that the Nyström-window acceleration preserves numerical fidelity across diverse perturbation regimes.

**Table 3:** Approximation accuracy of nyswin across synthetic scenarios. Mean relative error remains below 0.2%, substantially smaller than intrinsic permutation-based normalization variability.

| Scenario | dES (orig) | dES (nyswin) | Rel. Error (%) |
| --- | --- | --- | --- |
| Broad-spectrum | 0.2869 | 0.2872 | 0.108 |
| Strong targeted | 0.2169 | 0.2171 | 0.108 |
| Dose (high) | 0.3086 | 0.3090 | 0.121 |
| Dose (mid) | 0.2767 | 0.2771 | 0.127 |
| Dose (low) | 0.1830 | 0.1832 | 0.146 |
| Mild | 0.1369 | 0.1371 | 0.146 |
| Batch shift | 0.2321 | 0.2324 | 0.149 |
| Heavy-tailed | 0.3274 | 0.3279 | 0.166 |
| Mean/Max | — | — | **0.134 / 0.166** |

**Figure 7:**
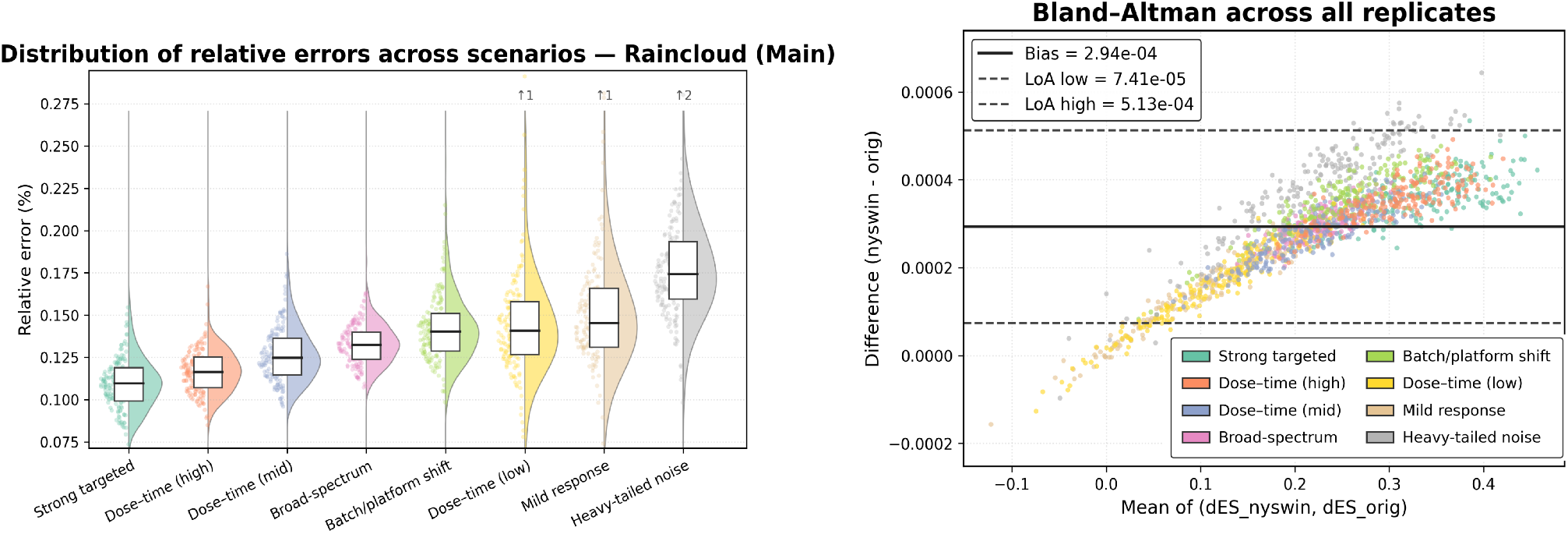
Approximation quality of nyswin. Comparison between exact dGSEA (`orig`) and the accelerated `nyswin` implementation across synthetic benchmarks. Left: scatter plot shows near-perfect agreement. Right: Bland–Altman plot confirms tight agreement with minimal bias. Numerical accuracy (Table 3) and distributional concordance demonstrate that `nyswin` preserves the quantitative semantics of dGSEA, rendering approximation error negligible for downstream inference and gradient-based optimization.

We evaluate computational performance on CPU and GPU backends as a function of gene number *G* (Fig. 8). Panel (a) shows GPU speedup of `nyswin` relative to the exact implementation. For small gene sets (*G* ≲ 3,000), approximation overhead and kernel launch costs dominate, limiting speedup. Beyond the crossover regime (*G* ≈ 3,378), reduced asymptotic complexity becomes dominant, yielding increasing gains that exceed 1.8× at *G* = 20,000. Panel (b) illustrates end-to-end runtime scaling: while classical GSEA remains efficient for single post hoc evaluations, the accelerated dGSEA implementation achieves practical GPU runtimes compatible with repeated training-time evaluation. Combined with broad robustness to hyperparameter variation (Appendix C), these results demonstrate that `nyswin` makes pathway-level objectives computationally feasible at genome scale.

**Figure 8:**
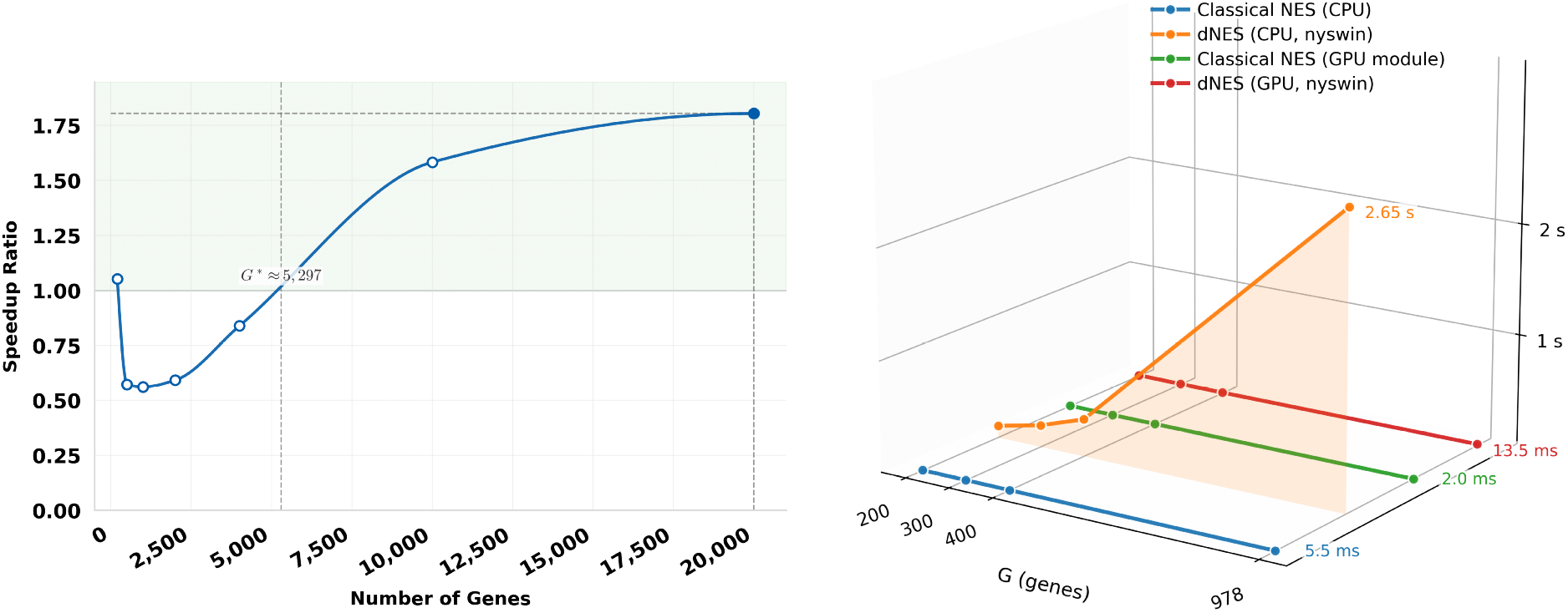
Computational performance of nyswin. (a) GPU speedup relative to exact implementation as a function of gene number *G*. Overhead dominates for small gene sets (*G <* 3,000); beyond the crossover at *G* ≈ 3,378, reduced asymptotic complexity yields increasing gains, exceeding 1.8 × speedup at *G* = 20,000. (b) End-to-end runtime scaling across CPU/GPU backends and classical/differentiable GSEA. While classical GSEA remains efficient for single evaluations, its non-differentiable nature precludes training-time integration. The accelerated dGSEA implementation achieves practical runtimes on GPU compatible with repeated evaluation, enabling its use as a structured objective within modern learning pipelines.

### 3.4 Pathway-aware training improves functional agreement without sacrificing gene-level fidelity

Having established that dGSEA preserves classical GSEA semantics with improved stability (Results 1–2) and can be evaluated efficiently at genome scale (Result 3), we now test its training-time utility as a structured supervision signal. We address a central question: can augmenting a standard gene-level objective with a pathway-level dGSEA term improve functional agreement quantified by pathway scores and enrichment directions without sacrificing gene-wise predictive fidelity? To isolate the effect of pathway-aware supervision, we employ a controlled ablation design where all model variants share the same backbone architecture and training protocol, differing only in the loss function.

#### Experimental design

We consider a SMILES-to-transcriptome prediction task using LINCS L1000 Level-5 signatures restricted to the 978 directly measured landmark genes. We retain compounds with valid canonical SMILES and at least three Level-5 instances, aggregating all profiles across cell line, dose, and duration by arithmetic averaging to form a single compound-level transcriptional signature. The final cohort contains *N* =10,554 compounds, split into train/validation/test subsets of sizes 7,916*/*1,055*/*1,583 using a fixed random seed. The pathway collection consists of the five curated directional pathways used in Result 2: p53 tumor suppressor, Cell cycle (CDK/Cyclin), Heat-shock (DNAJ/HSP40), Ubiquitin-proteasome (DUBs), and DNA damage (GADD45).

All model variants employ a frozen pretrained ChemBERTa encoder to map SMILES to molecular embeddings, followed by a lightweight regression head (Linear → ReLU+Dropout → Linear) that outputs a 978-dimensional transcriptional profile prediction **ŷ** ∈ ℝ^978^ (Fig. 9a). For numerical stability, we apply per-gene standardization during training: let ***µ, σ*** ∈ ℝ^978^ denote the per-gene mean and standard deviation estimated from the training set; gene-level losses are computed in standardized space **y**^(*z*)^ = (**y**^(orig)^ − ***µ***)*/****σ***, and predictions are mapped back to original space **ŷ**^(orig)^ = **ŷ**^(*z*)^⊙***σ***+***µ*** before pathway-level evaluation to avoid distortions introduced by standardization.

**Figure 9:**
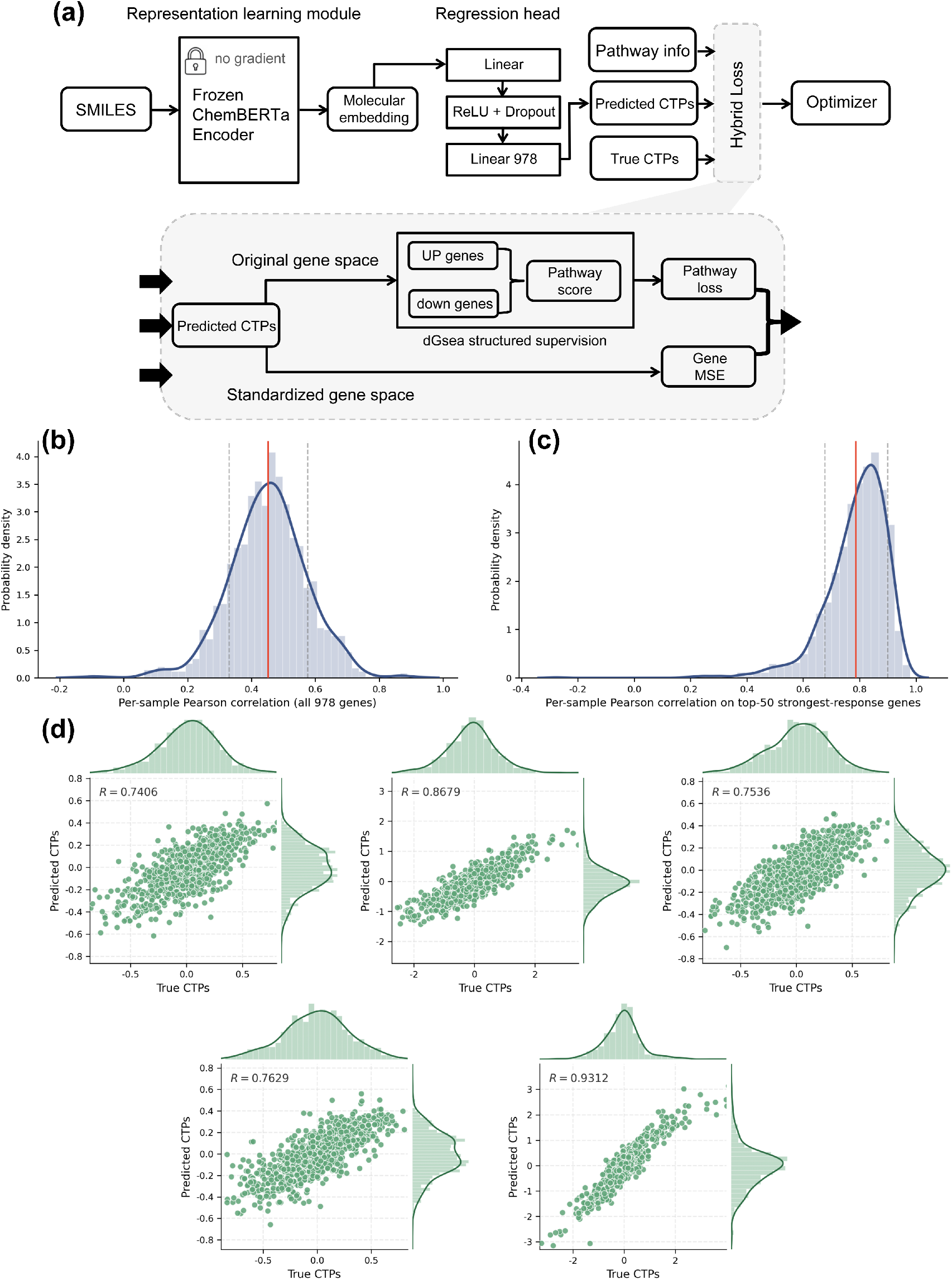
Pathway-aware training with dGSEA structured supervision. (a) Model architecture and loss overview: a frozen ChemBERTa encoder produces molecular embeddings, a regression head predicts CTPs, and training uses a hybrid objective combining gene-level losses (computed in standardized space) with dGSEA-based pathway supervision (computed in original space). (b) Distribution of per-sample Pearson correlations over all 978 genes on the test set shows that the hybrid objective preserves gene-level fidelity. (c) Distribution of per-sample correlations restricted to Top-50 strongest-response genes per sample demonstrates improved capture of dominant perturbation signals. (d) Representative predicted-versus-true scatter plots for selected test compounds illustrate gene-wise agreement across the full 978-dimensional profile.

#### Training objectives

We compare three loss configurations. The *baseline* (gene-level only) combines weighted MSE and correlation alignment in standardized space:

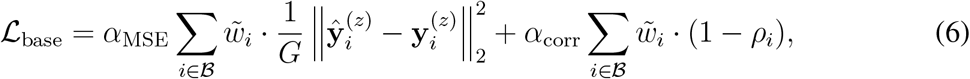

where 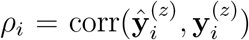 is the per-sample Pearson correlation and 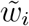 are normalized sample weights (based on replicate count). The *dGSEA-only* objective optimizes pathway-level supervision alone without gene-level constraints, serving as an ablation to assess whether pathway signals can be recovered without explicit gene-wise supervision. The *hybrid* objective combines both levels:

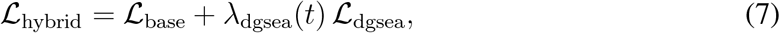

where *λ*_dgsea_(*t*) is a warm-up coefficient that increases with training iteration.

For each pathway *k* with UP set *U*_*k*_ and optional DOWN set *D*_*k*_, we define the signed pathway score via the differentiable enrichment operator:

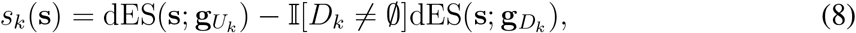

where 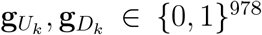 are indicator masks. Applying *s*_*k*_(·) to predictions and ground truth over a mini-batch yields pathway-score vectors **a** = *s*_*k*_(**Ŷ**^(orig)^) and **b** = *s*_*k*_(**Y**^(orig)^). The pathway loss penalizes mismatch using a composite objective:

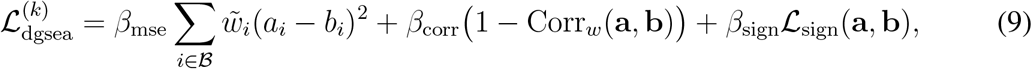

where Corr_*w*_ is weighted Pearson correlation, 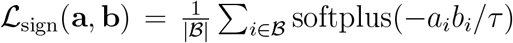 is a smooth sign-consistency penalty, and 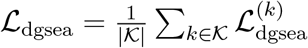 averages over all pathways.

#### Evaluation metrics

We evaluate both gene-level fidelity and pathway-level functional agreement. Gene-level performance is measured by per-sample Pearson correlation and RMSE between predicted and true transcriptional profiles over all 978 genes. To emphasize strong-response signals that dominate downstream interpretation, we additionally compute the same metrics on the Top-50 strongest-response genes per sample (selected by largest absolute ground-truth expression). Pathway-level performance is evaluated by mapping predicted and true profiles to pathway scores using the operator in Eq. (8) and reporting macro-averaged correlation, MSE, and sign accuracy across pathways. We additionally report *r*_95_, the 95th percentile of the per-sample Pearson correlation distribution on the test set.

#### Results

Tables 4 and 5, along with Fig. 9, summarize performance under the three objectives. The hybrid objective preserves gene-level fidelity matching or slightly improving the baseline in mean correlation (all genes: 0.449 → 0.452; 95th percentile: 0.651 → 0.656; Top-50: 0.786 → 0.788) and RMSE (all genes: 0.420 → 0.418; Top-50: 1.016 → 1.011) while substantially improving pathway-level functional agreement. Macro pathway correlation increases from 0.257 to 0.306 (+19%), sign accuracy from 0.620 to 0.641 (+3.4%), and pathway MSE decreases from 1.784 to 1.610 (-9.8%). Per-sample correlation distributions (Fig. 9b–c) confirm that the hybrid objective preserves gene-level performance, with Top-50 correlations showing strong capture of dominant perturbation signals. Representative scatter plots (Fig. 9d) illustrate tight gene-wise agreement for selected test compounds.

**Table 4:** Gene-level performance under controlled ablation. The hybrid objective preserves gene-level fidelity (mean *r* and RMSE comparable to baseline).

| Objective | Mean $r$ | $r_{95}$ | RMSE | Top-50 $r$ | Top-50 RMSE |
| --- | --- | --- | --- | --- | --- |
| Baseline | 0.449 | 0.651 | 0.420 | 0.786 | 1.016 |
| dGSEA-only | 0.292 | 0.463 | 30.467 | 0.633 | 34.234 |
| Hybrid (ours) | **0.452** | **0.656** | **0.418** | **0.788** | **1.011** |

**Table 5:** Pathway-level performance under controlled ablation (macro-averaged across pathways). The hybrid objective improves functional agreement (correlation, MSE, and sign accuracy) relative to the gene-level baseline.

| Objective | Correlation | MSE | Sign Accuracy |
| --- | --- | --- | --- |
| Baseline | 0.257 | 1.784 | 0.620 |
| dGSEA-only | 0.077 | 6.138 | 0.597 |
| Hybrid (ours) | **0.306** | **1.610** | **0.641** |

**Table 6:**
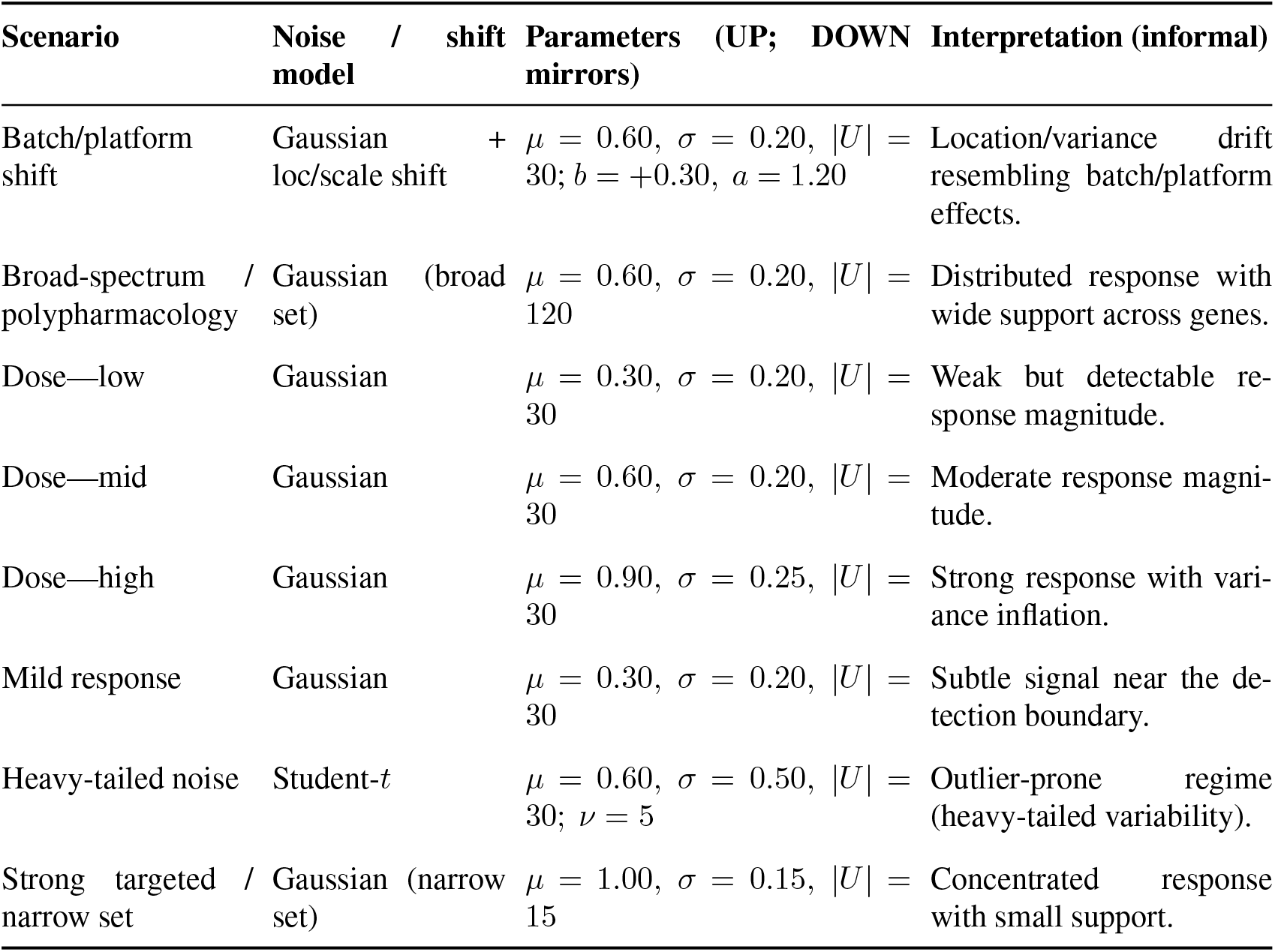
Controlled synthetic scenarios used for semantic validation.

In contrast, the dGSEA-only objective yields catastrophic gene-space reconstruction failure (mean *r* = 0.292, RMSE = 30.467 for all genes; Top-50 RMSE = 34.234), confirming that pathway-level supervision constrains set-level functionals but does not uniquely identify individual gene values. Notably, despite this gene-level collapse, dGSEA-only achieves limited pathway-level coherence (correlation = 0.077, sign accuracy = 0.597), demonstrating that pathway supervision can capture some directional enrichment patterns even without faithful transcriptional reconstruction. These results establish that dGSEA is most effective as an auxiliary structured objective rather than a standalone replacement for gene-level supervision: the hybrid formulation guides models toward decision-relevant, functionally coherent predictions while preserving the gene-level accuracy necessary for faithful transcriptional profiling.

## 4 Methods

We denote by *s* ∈ ℝ^*G*^ a real-valued score vector over *G* genes (e.g., a signed differential expression statistic), and by *g* ∈ {0, 1}^*G*^ the membership indicator of a gene set 𝒮 = {*i* : *g*_*i*_ = 1}; notation follows Appendix A. Unless stated otherwise, all vectors are column vectors and ranking is in *descending* order of *s*. This section first reviews the classical Gene Set Enrichment Analysis (GSEA) formulation (§4.1) and then introduces our new differentiable GSEA (dGSEA) statistic (§4.2).

### 4.1 Classical GSEA Formulation

We assume throughout that gene sets are non-trivial (1 ≤ |𝒮| ≤ *G* − 1) and that the total weighted score of genes within the set is non-zero, ensuring that denominators in the formulation are well-defined.

Given a score vector *s* and a gene set indicator *g*, let *π* be the permutation that sorts *s* in descending order, i.e., *s*_*π*(1)_ ≥ · · · ≥ *s*_*π*(*G*)_. For a power parameter *p* ≥ 0, the hit and miss weights for each gene in the sorted list are defined as:

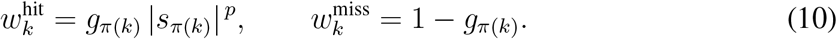

Let 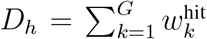 and 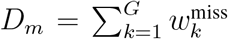 (note that *D*_*m*_ = |𝒮^∁^| for *p* = 0). The classical running-sum curve is constructed by accumulating these weights:

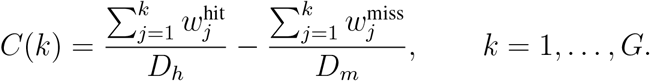

The Enrichment Score (ES) is the maximum deviation of this curve from zero, formally defined as:

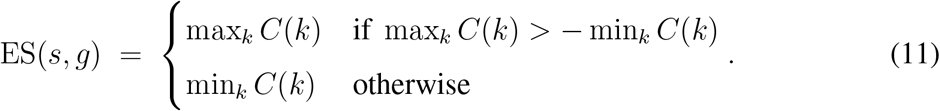

To account for gene set size and score distribution, a permutation-based null distribution is generated to yield a Normalized Enrichment Score (NES). One computes ES scores for *K* permutations of the gene labels, ES^(*k*)^, to estimate the conditional expectations of the null distribution:

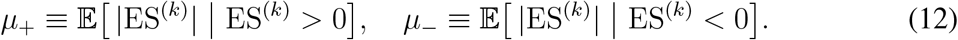

The NES is then the observed ES normalized by the expectation corresponding to its sign: NES = ES*/µ*_*±*_.

### 4.2 Differentiable GSEA (dGSEA)

The classical GSEA statistic is non-differentiable with respect to the input scores *s* due to the hard ranking operation and the stepwise nature of the running sum. To overcome this limitation, we introduce smooth, differentiable relaxations for each component of the GSEA pipeline.

#### (i) Soft Ranking

To replace the hard ranking permutation *π*, we define a differentiable soft rank. Given a temperature parameter *τ*_rank_ *>* 0, we first compute a pairwise soft-comparison matrix using the sigmoid function *σ*(*z*) = (1 + *e*^−*z*^)^−1^:

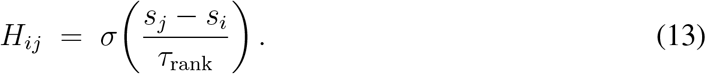

*H*_*ij*_ approximates whether gene *j* has a higher score than gene *i*. The soft rank *r*_*i*_ for gene *i* is then formulated as:

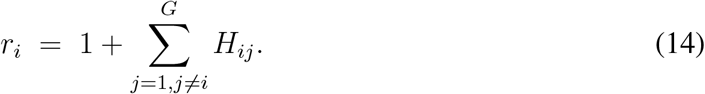

#### (ii) Smooth Prefix Accumulation

Next, we replace the discrete running sum with a smooth prefix accumulation. For a temperature *τ*_prefix_ *>* 0, we define a smooth prefix indicator *H*_*i*_(*t*), which approximates whether gene *i* falls within the top *t* ranks:

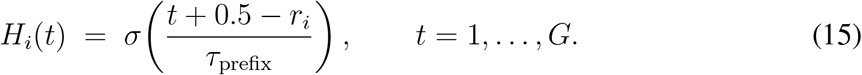

Using this indicator, we define the smoothed cumulative hit and miss curves, 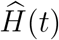 and 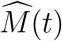, over the entire range of ranks:

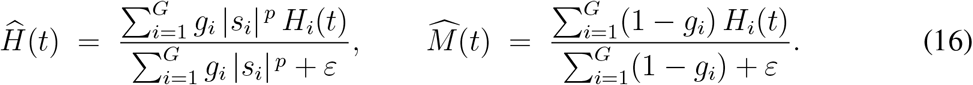

The difference between these curves yields a smooth, differentiable running-sum curve, 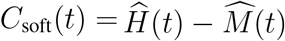.

#### (iii) Differentiable Enrichment Score Aggregator

To find the maximum deviation of *C*_soft_(*t*) in a differentiable manner, we employ a temperature-controlled softmax aggregation. Given a temperature *τ*_abs_ *>* 0, the weights *ω*_*t*_ are computed based on the magnitude of the deviation at each position:

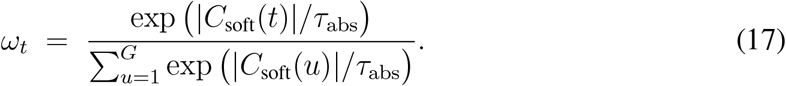

The differentiable Enrichment Score, dES, is the weighted sum of the signed deviations:

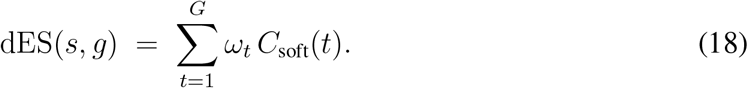

This formulation is fully differentiable with respect to the input scores *s*.

#### (iv) Normalization and Statistical Significance (dNES)

To ensure comparability across different gene sets and experiments, we define a normalized score, dNES, mirroring the statistical logic of the classical NES. This involves a robust permutation-based normalization and a calibration step.

##### Sign-specific robust denominator

We generate a null distribution by computing dES^(*k*)^ over *K* permutations. The normalization denominators are estimated using a robust procedure combining a trimmed mean and a Winsorized mean. For a trimming proportion *ρ* ∈ (0, 1*/*2) and shrinkage coefficient *λ* ∈ [0, 1], the estimator 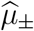 is defined as 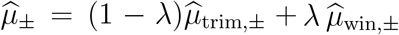.

##### κ–calibration to classical scale

To enhance interpretability, we introduce a calibration factor, *κ*, that aligns the scale of the dGSEA null distribution with the classical one. We compute robust means for both the classical null 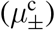 and the differentiable null 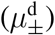 from the same permutations, and define the factor as their ratio:

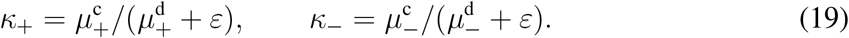

The final calibrated denominator is 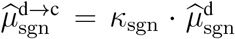 . The final reported score is the fully calibrated dNES:

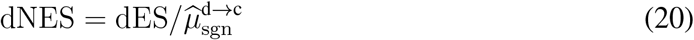

Significance is determined via one-sided *p*-values derived from the empirical permutation-based null distribution.

#### Practical hyperparameter setting

The differentiable formulation introduces three temperature parameters, (*τ*_rank_, *τ*_prefix_, *τ*_abs_), which control soft ranking, smooth prefix accumulation, and extremum aggregation, respectively. Unless otherwise specified, we use the default setting (0.80, 1.10, 0.70) throughout the experiments.

These values were selected based on a comprehensive sensitivity analysis reported in Appendix C. As shown there, dGSEA exhibits a broad plateau of high agreement with classical GSEA across moderate parameter variations, with Spearman correlations exceeding 0.9 within *τ*_rank_ ∈ [0.80, 1.10] and *τ*_prefix_ ∈ [0.80, 1.20]. Among the three parameters, *τ*_abs_ primarily controls extremum sharpness, while *τ*_rank_ and *τ*_prefix_ demonstrate low sensitivity within practical ranges. These results indicate that dGSEA is robust to moderate hyperparameter perturbations, and extensive tuning is not required for stable behavior.

#### Algorithmic formulation

Algorithm 1 summarizes the core differentiable enrichment operator that maps a gene-level score vector to a scalar dES. This formulation is free of acceleration or statistical post-processing and serves as the conceptual anchor for subsequent algorithms.

##### Algorithm 1

Differentiable enrichment score (dES)

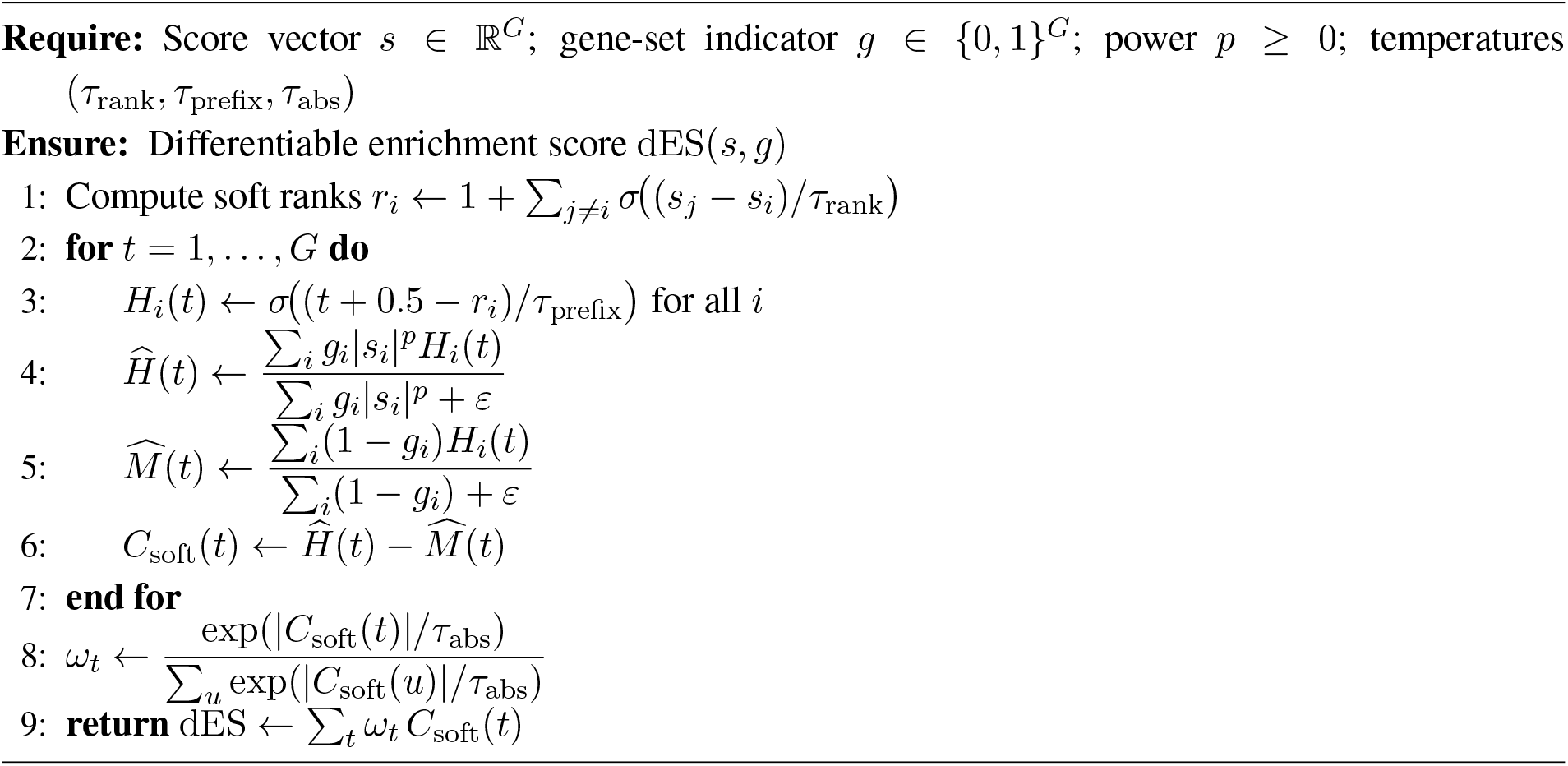

Algorithm 2 describes the statistical post-processing that converts dES into a normalized score (dNES) and a permutation-based *p*-value. This step is independent of any acceleration strategy.

##### Algorithm 2

Sign-specific normalization and permutation inference for dGSEA

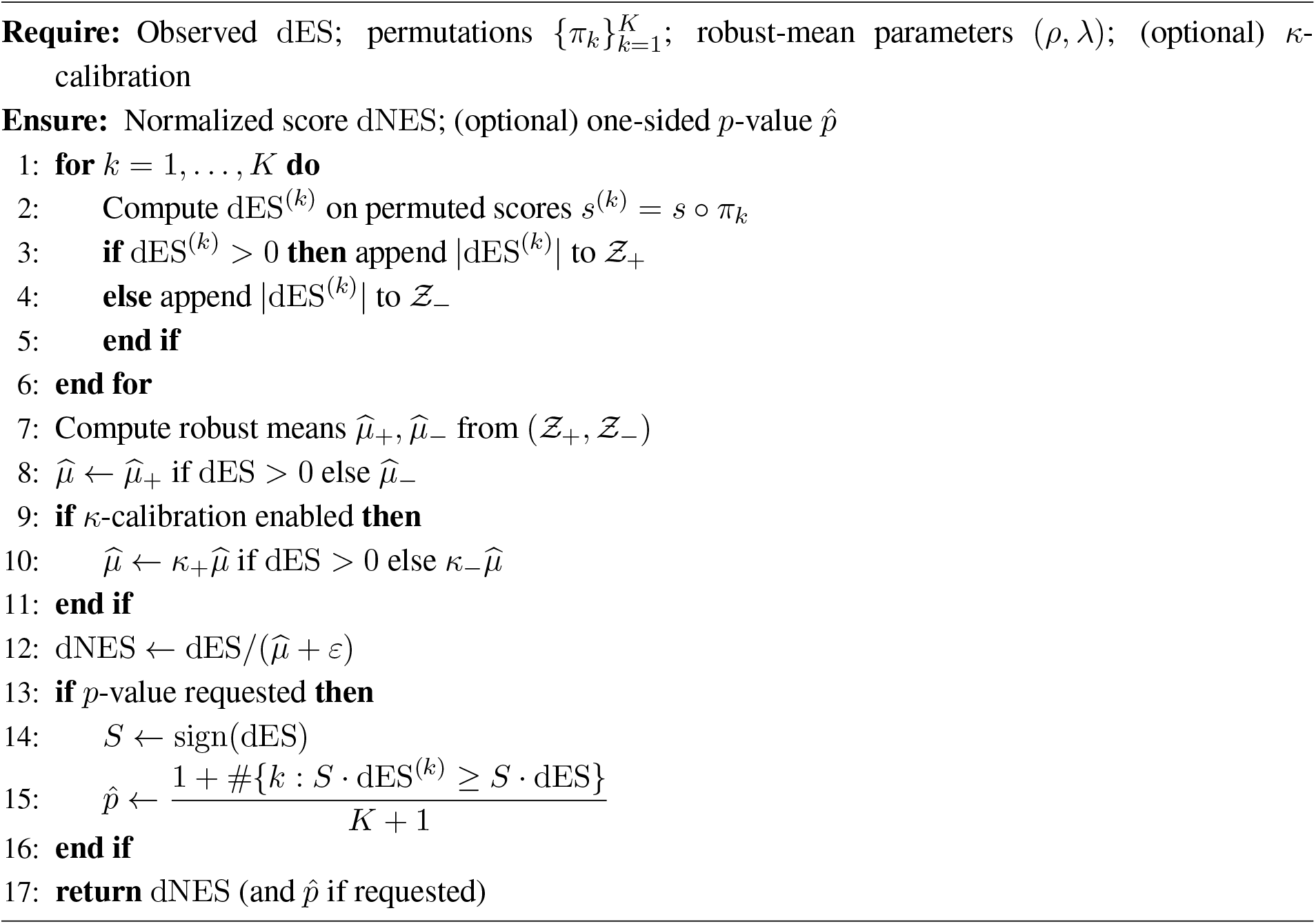

### 4.3 Scalable Implementation and Performance Optimization

The dGSEA formulation presented in §4.2, while theoretically elegant, faces severe computational challenges in its naive implementation. Its quadratic complexity makes it impractical for genome-scale analysis. For instance, on a typical L1000 gene expression profile with 12,328 genes, a single score evaluation using the highly optimized classical GSEA implementation takes less than a second, whereas our naive dGSEA requires over two minutes. This performance gap renders it unusable for large-scale applications, such as model training or screening vast libraries of gene sets.

This section details a series of algorithmic and engineering optimizations designed to overcome these bottlenecks, transforming dGSEA into a high-performance, scalable tool. To provide theoretical guarantees for these optimizations, we will also present and prove several key theorems and propositions.

#### 4.3.1 Analysis of Computational Bottlenecks

The prohibitive cost of the naive dGSEA formulation originates from two core components, both of which exhibit quadratic complexity with respect to the number of genes, *G*:

1. **Quadratic Complexity of Soft Ranking:** The soft rank calculation in Eq. (14) requires an all-pairs comparison between genes, resulting in a time and memory complexity of *O*(*G*^2^). For a typical human gene panel (*G* ≈ 20, 000), this would involve approximately 4 × 10^8^ operations, consuming hundreds of gigabytes of memory.
2. **Quadratic Complexity of Prefix Accumulation:** Even with pre-computed ranks, calculating the smooth running-sum curve *C*_soft_(*t*) across the full grid of ranks (*t* = 1, …, *G*) requires evaluating the prefix kernel *H*_*i*_(*t*) for each gene at each position, leading to an overall complexity of *O*(*G*^2^).

#### 4.3.2 Algorithmic Acceleration Strategies

To address these bottlenecks, we introduce two algorithmic acceleration strategies that dramatically reduce the computational workload while maintaining theoretical guarantees on accuracy.

##### Accelerating Soft Ranks via Nyström Approximation

To break the *O*(*G*^2^) barrier of soft ranking, we employ the Nyström method [28] to approximate the all-pairs comparison matrix. The core idea is to select a small, representative subset of *m* ≪ *G* “anchor points” from the score distribution. The rank of each gene is then approximated by comparing it only to these *m* anchors. The Nyström-approximated soft rank, 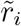, is given by:

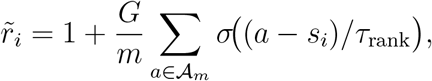

where 𝒜_*m*_ is the set of *m* anchors, typically chosen as quantiles of the score vector *s*. This reduces the complexity of this step to *O*(*Gm*). The accuracy of this approximation is formally guaranteed by the following robust theorem.

*Remark* 4.1 (Star-discrepancy of empirical quantile anchors). If 𝒜_*m*_ are the *m* equally spaced empirical quantiles of *F*_*G*_, then ∥*F*_*A*_ − *F*_*G*_∥_∞_ ≤ 1*/m* + *O*(1*/G*); in particular ∥*F*_*A*_ − *F*_*G*_∥_∞_ = *O*(1*/m*) when *m* ≪ *G*.

##### Accelerating Prefix Accumulation with a Windowed Grid

Based on the domain knowledge that the ES extremum typically occurs at the tails of the ranked list, we restrict the computation of *C*_soft_(*t*) to a “windowed grid,” 𝒯 . This reduces the prefix accumulation complexity from *O*(*G*^2^) to *O*(*GT* ). The following proposition provides a formal error bound for this strategy and establishes the conditions under which the error vanishes.

Algorithm 3 implements an accelerated approximation of Algorithm 1 by replacing full soft ranking and prefix evaluation with Nyström anchors and a windowed grid.

###### Algorithm 3

`nyswin`: Nyström–Window accelerated dES

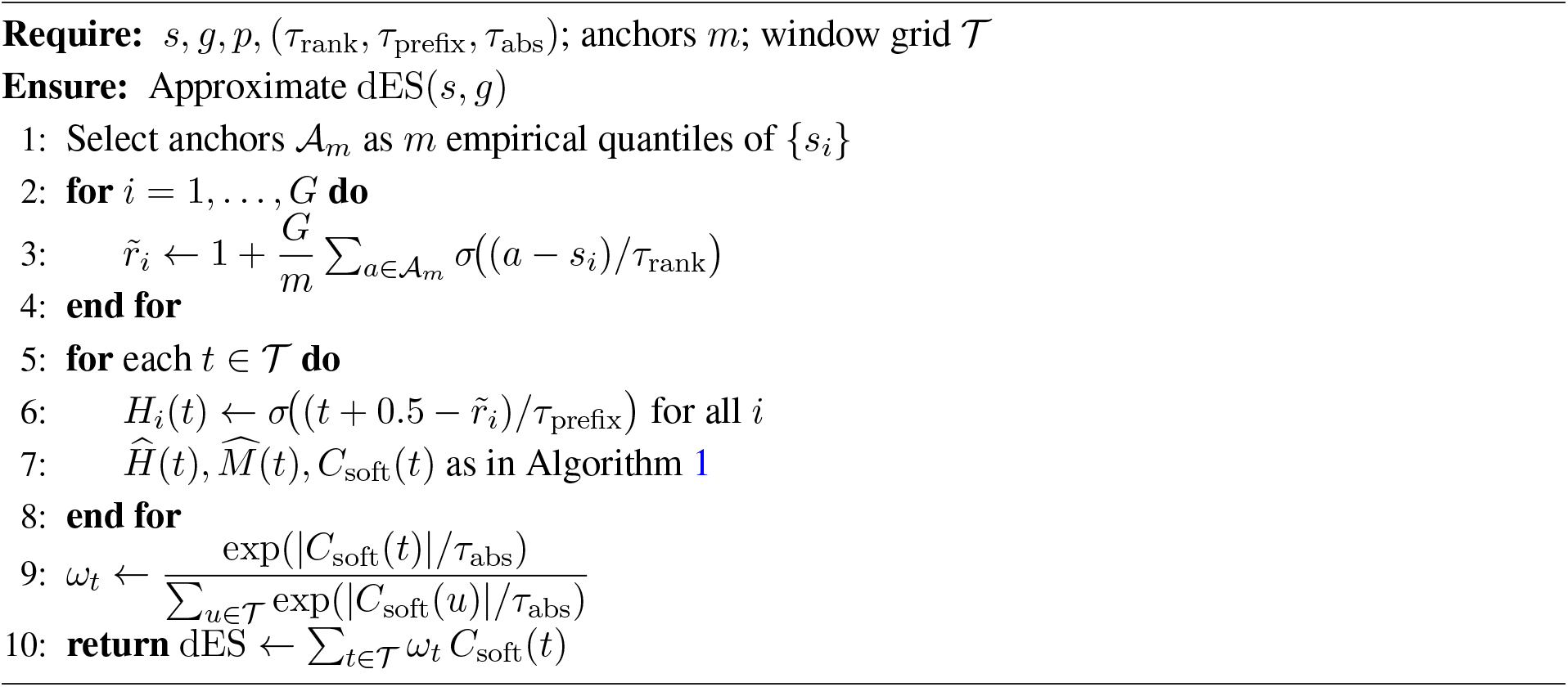

#### 4.3.3 Engineering and Hardware Acceleration

The aforementioned algorithms are implemented with a focus on maximizing throughput on modern parallel hardware, particularly GPUs. All core computations are expressed as batched tensor operations. Key engineering optimizations include the use of **shared permutations** across all gene sets in a batch to minimize redundant computations, and a **chunking** strategy to process a large number of permutations (*K*) in smaller slices, thereby controlling the peak GPU memory footprint.

#### 4.3.4 Summary of the Accelerated Algorithm and Performance Analysis

The combination of the Nyström approximation and the windowed grid results in a highly efficient algorithm, which we term the `nyswin` variant. This variant eliminates the quadratic complexity bottlenecks of the naive formulation. The performance of all variants is formally summarized by the following theorem.

##### Practical Guidance

For any application involving genome-scale data (*G* ≳ 1, 000), the `nyswin` variant is strongly recommended. The `orig` variant, while being the direct implementation of the core theory, serves primarily as a theoretical baseline and is only practical for very small gene universes. A detailed comparison of runtimes on empirical data is provided in the Experiments section.

## 5 Discussion

### From gene-level objectives to functional-level supervision

A common setup in transcriptomic prediction is to optimize upstream models with gene-wise objectives (e.g., MSE or correlation), whereas downstream interpretation and screening often operate on gene sets via rank- and set-dependent statistics such as enrichment, connectivity, or reversal scoring. Prior work has noted that strong gene-level metrics do not necessarily translate into stable pathway-level conclusions; when profile prediction is imperfect, modest but systematic ranking errors can change enrichment direction or reorder pathways (Fig. 1). We take this objective–functional mismatch as the motivating premise of the present study and focus on how to make downstream, decision-relevant set-level criteria usable during training.

Here we use *set-level functional* to mean a mapping from a gene score vector (e.g., predicted differential expression) and a gene-set definition to a scalar pathway score (as in classical GSEA). Such functionals are typically applied post hoc because their native forms are non-differentiable (due to hard ranking and extremum selection) and can be costly to evaluate repeatedly. Our goal is therefore practical: to construct a differentiable, calibrated, and scalable surrogate that preserves the intended semantics closely enough to serve as a training-time signal. Instantiating this with GSEA, dGSEA provides a smooth surrogate with a calibrated normalization (dNES) and an accelerated implementation (nyswin), enabling integration into gradient-based optimization. In E3, using dGSEA as an auxiliary structured term improves pathway-level agreement (correlation and sign consistency) while leaving gene-level accuracy essentially unchanged under our fixed SMILES-to-CTPs backbone; in contrast, optimizing the pathway term alone degrades gene-level reconstruction. This pattern supports treating dGSEA as a complementary constraint akin to structured regularization rather than as a replacement for gene-wise supervision.

### A general construction paradigm: soften, align, and accelerate

Although we instantiate our approach with GSEA, the design reflects a broader recipe for turning discrete, rank-based set-level criteria into training-time objectives. The recipe couples: (i) *softening* non-smooth operators (ranking, prefix indicators, extremum selection) via differentiable relaxations (Eq. (4– 9)) with limit consistency and controlled aggregation error (Theorem 4.1; Proposition 4.2); *alignment* of the surrogate to the classical scale through robust, sign-specific permutation normalization and *κ*-calibration (Eq. (10)), preserving the null semantics and permutation validity (Theorem 4.3); and (iii) *acceleration* that makes repeated evaluation feasible by reducing the dominant quadratic terms using Nyström approximation and a windowed prefix grid (Theorem 5.3). Together, these components provide a practical blueprint whenever downstream evaluation depends on rank-sensitive, set-structured functionals while upstream learning relies on smooth losses.

### Limitations and future work

The present study has several limitations that delimit the scope of its claims.

First, our evaluation is computational and statistical rather than experimental. While we show that dGSEA improves agreement with classical GSEA and enhances pathway-level fidelity under structured supervision, we do not assess whether pathway-consistent predictions translate into improved biological outcomes in downstream assays. Accordingly, our results should be interpreted as addressing an algorithmic and statistical gap between upstream training objectives and downstream functional analyses, rather than establishing causal biological efficacy. An important next step is to evaluate pathway-aware training in prospective screening or mechanism-of-action studies with experimental validation.

Second, the pathway collections used in E2 and E3 are intentionally small and curated to enable controlled analysis and interpretability. Although this setting isolates the effect of structured supervision, it does not address behavior under large, redundant, and highly overlapping pathway libraries. Extending pathway-aware supervision to broader gene set collections raises statistical and optimization questions, including the treatment of pathway overlap and differential pathway reliability, which remain open.

Third, dGSEA inherits the assumptions and limitations of gene-set-based representations of biology. If pathway annotations are incomplete, context-inappropriate, or poorly specified for a given assay, the induced supervision may bias learning toward an imperfect prior. Pathway-aware objectives should therefore be viewed as a mechanism for injecting interpretable biological structure, not as a replacement for careful pathway curation.

Beyond these limitations, a notable advantage of dGSEA is its modularity. The method is agnostic to model architecture and can be integrated into context-aware predictors that account for cell type, dose, or time, rather than being restricted to structure-only settings. As future work, we plan to release pretrained models together with an open fine-tuning framework that enables pathway-aware adaptation of existing predictors by specifying downstream pathway collections, facilitating controlled evaluation of functional-level objectives in diverse application settings.

## 6 Conclusion

In this work, we address a fundamental but often overlooked disconnect in transcriptomic prediction pipelines: models are trained with gene-level objectives, whereas scientific interpretation and decision-making are performed at the level of pathways and gene sets. When predictive accuracy is limited, this objective mismatch can lead to unstable or misleading functional conclusions, even when conventional gene-wise metrics appear strong.

We propose dGSEA as a practical resolution to this mismatch. By constructing a differentiable, calibrated, and scalable surrogate of classical GSEA, dGSEA enables pathway-level enrichment statistics to be used directly as training-time supervision. Our analysis establishes that dGSEA preserves the essential semantics of classical GSEA, including rank-based behavior, permutation validity, and normalized score interpretation, while remaining compatible with gradient-based optimization. Through extensive synthetic and real-data experiments, we show that dGSEA closely tracks classical GSEA, exhibits improved stability under noise, and serves as an effective structured regularizer that improves pathway-level coherence without degrading gene-level predictive performance.

Beyond its specific instantiation for GSEA, dGSEA exemplifies a broader paradigm for integrating downstream, decision-relevant functionals into upstream learning. By combining differentiable relaxation, statistical calibration, and algorithmic acceleration, this paradigm provides a general blueprint for aligning training objectives with the functional criteria that ultimately guide scientific inference. We expect this approach to be broadly applicable to other rank- and set-based analyses in computational biology and beyond.

## Acknowledgements

This work was supported by JSPS KAKENHI Grant Number JP23H03356.

## Code availability

An open-source implementation of dGSEA is available at https://github.com/LeeShuaiyu/dgsea-paper-code.

## A Experimental protocols supporting Results

This appendix records the experimental setups that support the Results section, focusing on data generation/preparation, permutation protocols, and evaluation configurations. To improve readability, we present the protocols in two parts: a controlled benchmark for semantic validation (Appendix A.1) and a real-data enrichment pipeline on LINCS L1000 (Appendix A.2).

### Permutation convention

Whenever enrichment is evaluated for a single gene-score vector *s* ∈ ℝ^*G*^, permutations act on gene indices (gene-label permutations): for a permutation 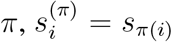. All cross-method comparisons reuse *shared permutation indices* to ensure paired null samples.

### A.1 Controlled benchmark for semantic validation

We generate synthetic score vectors to isolate how enrichment behavior changes under controlled signal strength, support size, heavy-tailed noise, and global location/scale drift. Unless otherwise stated, we use *G* = 400 genes. For each instance we sample disjoint UP and DOWN index sets *U, D* ⊂ {1, …, *G*} with |*U* | = |*D*| and define indicator vectors *g*_up_, *g*_dn_ ∈ {0, 1}^*G*^ by membership. Scores are generated by injecting a signed mean shift on *U* and *D* with additive background noise:

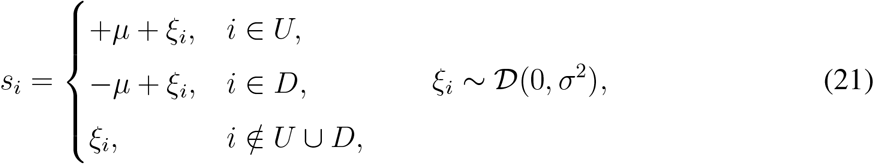

where 𝒟 is scenario-dependent: Gaussian or Student-*t* with degrees-of-freedom *ν*. For Student-*t*, the scale is set so that the effective noise magnitude is comparable under the specified (*σ, ν*). To emulate platform/batch drift, the batch/platform-shift scenario applies a global location/scale transform after sampling (21):

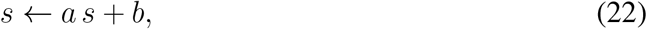

with fixed (*a, b*).

We use eight scenarios spanning weak/strong signals, narrow/broad support, heavy-tailed noise, and location/variance drift. Parameters are reported for the UP set; the DOWN set mirrors the UP set. *Interpretations* are informal labels of synthetic regimes and do not constitute mechanistic claims.

Classical GSEA uses the weighted running-sum formulation in Methods with power parameter *p* = 1 and average-rank tie handling. dGSEA uses temperatures (*τ*_rank_, *τ*_prefix_, *τ*_abs_) = (0.80, 1.10, 0.70) unless stated otherwise. dNES is computed using sign-specific robust normalization (trimmed/Winsorized robust mean) and optionally *κ*-calibrated to the classical NES scale. All permutation nulls use gene-label permutations (Appendix A), and permutation indices are shared across methods for paired comparison.

Controlled evaluations use the following fixed settings: (i) score agreement uses *M* = 32 independent instances with *K* = 256 shared permutations; (ii) null calibration uses *M* = 120 null instances with *K* = 200 permutations each, summarized by QQ plots and uniformity tests (KS and Berk–Jones), with BH-FDR illustrated at *α* = 0.05; (iii) the opposing-signal reversal test repeats 50 runs and reports mean±SEM; (iv) additive-noise robustness adds 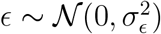 with *σ*_*ϵ*_ ∈ {0.2, 0.5, 1.0} and repeats each condition 40 times, summarizing instability by Std(ΔScore) relative to the noiseless baseline; and (v) hyperparameter sensitivity sweeps *τ*_rank_ ∈ [0.8, 1.6], *τ*_prefix_ ∈ [0.8, 1.2], *τ*_abs_ ∈ {0.40, 0.60, 0.80, 1.00}, reporting Spearman(dNES, NES). When benchmarking approximation fidelity and scalability, we compare `orig` and `nyswin` under identical inputs and shared permutations; unless stated otherwise, `nyswin` uses Nyström anchors *m* = 256 and permutation chunk size *c* = 8.

### A.2 Real-data enrichment pipeline on LINCS L1000

We use LINCS L1000 (GEO: GSE92742), Level-5 signatures, restricted to the 978 directly measured landmark genes. We retain small-molecule perturbations with valid canonical SMILES, enforce molecular validity via RDKit parsing, and exclude SMILES with fewer than three Level-5 instances to stabilize compound-level signatures. For each canonical SMILES, we aggregate all corresponding Level-5 profiles across cell line, dose, and duration by arithmetic averaging, producing one compound-level signature in ℝ^978^. We record `sample_count` (number of aggregated instances) and `unique_cells` (number of unique cell lines) per compound. The enrichment concordance cohort contains 10,554 compound-level signatures.

We evaluate five curated directional pathways (UP set + optional DOWN set), mapped to the 978-gene space. HeatShock_DNAJ intentionally includes one missing gene (DNAJB2) to test robustness under incomplete coverage.

For each compound and pathway, we compute classical NES and dGSEA dNES using gene-label permutations with *K* = 300 and fixed random seed 7. dGSEA uses the accelerated `nyswin` implementation with Nyström anchors *m* = 256 and permutation chunk size *c* = 8, and temperatures (*τ*_rank_, *τ*_prefix_, *τ*_abs_) = (0.80, 1.10, 0.70).

Directional regulation is summarized by

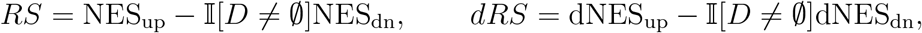

with *RS* = NES_up_ (and similarly *dRS*) for pathways without a DOWN component. Agreement between *RS* and *dRS* across compounds is measured by Spearman correlation, Kendall’s *τ*, and concordance index (C-index). We also analyze absolute rank differences and Top-*K* / Top-50 overlap. To assess whether method-specific Top-50 subsets exhibit non-random chemical structure, we compute Morgan fingerprints for the Top-50 union set and evaluate intra-group homogeneity by mean pairwise Jaccard similarity, with significance assessed by a 10,000-resample permutation test under the null that group membership is random given group sizes. t-SNE embeddings are used for visualization only.

## B Theoretical guarantees and proofs

This appendix provides formal statements and proofs for the theoretical results cited in Results 2–3 and Methods §4.2. We use the same notation as in the main text. All permutations in *p*-value arguments are gene-label permutations on *s* ∈ ℝ^*G*^.

### B.1 Asymptotic Properties and Consistency

We first establish that dGSEA converges to the classical GSEA statistic as its smoothing parameters approach zero.

#### Theorem B.1

(Limit Consistency, dES → ES). *Let the temperature parameters τ*_rank_, *τ*_prefix_, *τ*_abs_ → 0, *and let the prefix grid be the full set of ranks* 𝒯 = {1, …, *G*}. *Then, the differentiable enrichment score converges to the classical enrichment score:*

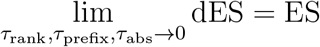

*Proof (in the distinct-scores, unique-argmax case)*. **Step 1 (soft-rank** → **hard rank)**. Let *i* be fixed. If *s*_*j*_ *> s*_*i*_, then (*s*_*j*_ − *s*_*i*_)*/τ*_rank_ → +∞, which implies *σ*((*s*_*j*_ − *s*_*i*_)*/τ*_rank_) → 1. If *s*_*j*_ *< s*_*i*_, the limit is 0. Therefore,

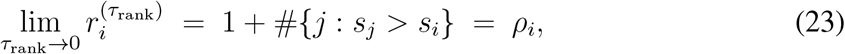

where *ρ*_*i*_ is the standard descending rank.

**Step 2 (prefix kernel** → **indicator function)**. For a fixed integer *t*, as 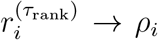 and letting *τ*_prefix_ → 0,

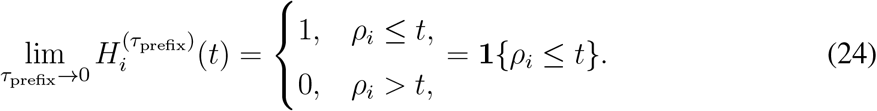

This holds because for distinct scores *ρ*_*i*_ ∈ ℤ and hence *t* + 0.5 − *ρ*_*i*_ ≠ 0, so the sigmoid does not converge to its threshold.

**Step 3 (***C*_**soft**_ → *C* **uniform convergence)**. Substituting the limit from Step 2 into the definition of 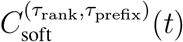, we obtain pointwise convergence for each *t*:

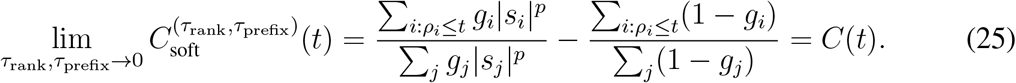

Since the convergence is governed by bounded functions on a finite set 𝒯, the convergence is uniform: sup_*t*∈*T*_ |*C*_soft_ − *C*| → 0.

**Step 4 (Softmax selects the extremum)**. Let *y*_*t*_ = |*C*(*t*)|, assume it has a unique maximum at *t*^⋆^, and let the gap be 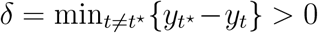. From the uniform convergence in Step 3, for any *ε < δ/*3, there exists *τ*_0_ *>* 0 such that for *τ*_rank_, *τ*_prefix_ ∈ (0, *τ*_0_), we have ||*C*_soft_(*t*)| − *y*_*t*_| ≤ *ε* for all *t*. This implies |*C*_soft_(*t*^⋆^)| − |*C*_soft_(*t*)| ≥ *δ* − 2*ε > δ/*3 for all *t* ≠ *t*^⋆^. Consequently, the softmax weight 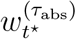 converges to 1 as *τ*_abs_ → 0, while all other weights converge to 0. It follows that:

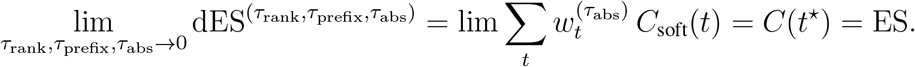

*Remark* B.1 (Handling Ties). If scores are not distinct, the proof remains valid by adopting either of two standard tie-handling schemes: (i) **Average Rank**: The classical rank *ρ*_*i*_ is defined as the average rank for ties. The limit of the soft rank 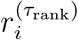 converges to this average rank, as 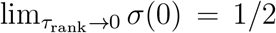. The rest of the proof follows analogously. (ii) **Vanishing Jitter**: A small random perturbation is added to the scores, *s*^(*ϵ*)^ = *s* + *η*, which breaks ties with probability 1. The theorem holds for any fixed *ϵ >* 0. By continuity, the limits as *ϵ* → 0 are consistent.

The following proposition provides a quantitative bound on the error introduced by the softmax aggregation.

#### Proposition B.2

(Error Bound for Softmax Aggregation (Corrected)). *Let* 𝒯 *be the prefix grid and define w*_*t*_ ∝ exp |*C*_*soft*_(*t*)|*/τ*_abs_ *with* ∑ _t∈*T*_ *w*_*t*_ = 1. *Let*

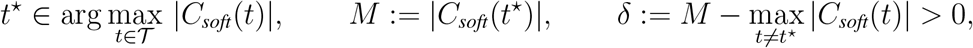

*and write s*^⋆^ := sign *C*_*soft*_(*t*^⋆^) . *Then the following bound holds:*

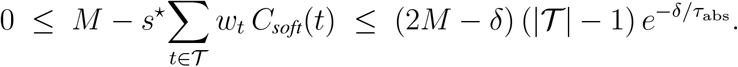

*Moreover, if all non-maximal points share the same sign as t*^⋆^, *i*.*e*. sign *C*_*soft*_(*t*) = *s*^⋆^ *for all t* ≠ *t*^⋆^, *then with* Δ_*t*_ := *M* − |*C*_*soft*_(*t*)| ≥ *δ*,

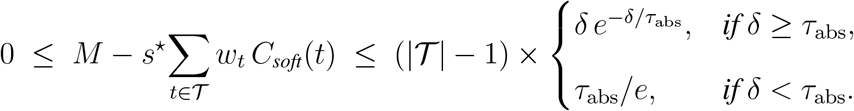

*Proof*. Fix *t*^⋆^ as above and set Δ_*t*_ := *M* − |*C*_soft_(*t*)| for *t* ≠ *t*^⋆^. Then Δ_*t*_ ≥ *δ* and

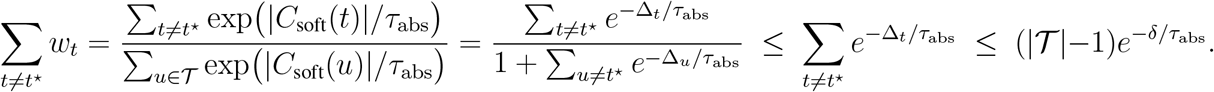

For the general bound,

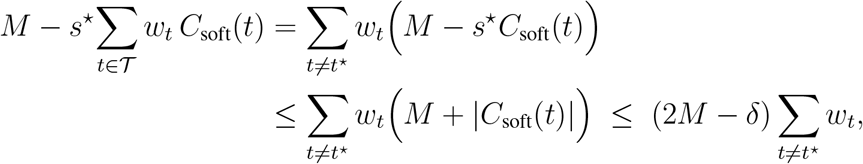

because |*C*_soft_(*t*)| ≤ *M* − *δ* for *t* ≠ *t*^⋆^. Combining with the weight bound yields

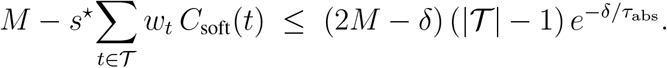

For the refined bound, assume sign(*C*_soft_(*t*)) = *s*^⋆^ for all *t* ≠ *t*^⋆^. Then

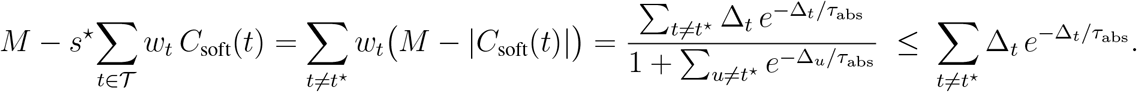

The function 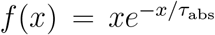 attains its maximum on [*δ*, ∞) at *x* = *δ* if *δ* ≥ *τ*_abs_, and at *x* = *τ*_abs_ with value *τ*_abs_*/e* if *δ < τ*_abs_. Therefore

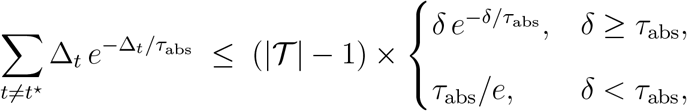

which proves the claim.

### B.2 Statistical Validity and Robustness

#### Theorem B.3

(Validity of Permutation *p*-values). *Let a one-sided p-value be computed from K permutations as*

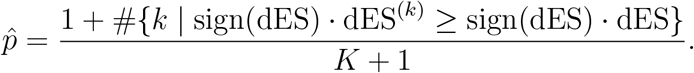

*Under the null hypothesis that the gene set membership is independent of the scores*, 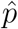 *provides valid control of the Type I error rate. That is, it is* ***super-uniform***: 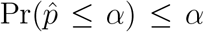 *for any α* ∈ [0, 1].

*Proof*. Let *U*_*i*_ := dES^(*i*)^ for *i* = 0, 1, …, *K* where *U*_0_ = dES is the observed value and *U*_1_, …, *U*_*K*_ are from i.i.d. uniform permutations under the null. Then (*U*_0_, …, *U*_*K*_) is exchangeable. Define *S* := sign(*U*_0_) and apply the same transformation to all components: *V*_*i*_ := *S* · *U*_*i*_. This map is either the identity (*S* = +1) or an order-reversing map applied to *all* coordinates (*S* = −1); in either case, the rank of *V*_0_ among 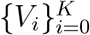 is still discrete-uniform on {1, …, *K* + 1}. Note that

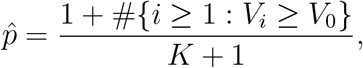

so 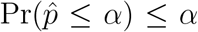 for any *α* ∈ [0, 1], i.e. 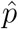 is super-uniform. If sample splitting is used, the same argument applies *conditionally* on the normalization subset, hence validity holds unconditionally.

#### Proposition B.4

(Consistency and Robustness of Normalization). *Fix a trimming proportion ρ* ∈ (0, 1*/*2) *and a shrinkage coefficient λ* ∈ [0, 1]. *For each sign* ±, *consider the permutation null samples Z*_*i,±*_ := |dES^(*i*)^| *restricted to* {sign(dES^(*i*)^) = ±}. *Let* 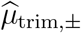 *and* 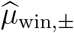 *be the ρ-trimmed and ρ-Winsorized means of* {*Z*_*i,±*_}, *respectively, and define the estimator as* 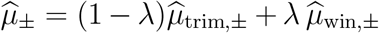. *If* 𝔼[*Z*_*i,±*_ ] *<* ∞, *then as the number of permutations K* → ∞, 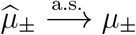, *its population counterpart. Moreover, the breakdown point of* 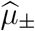 *equals ρ*.

*Proof*. Both 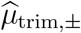 ^and 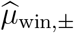^ are *L*-statistics; by the strong law for *L*-statistics, they converge almost surely to their population functionals. Slutsky’s theorem yields the almost sure convergence of their convex combination 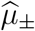. For robustness, under *ϵ*-contamination with *ϵ < ρ*, all outliers are trimmed or capped by construction, so the estimator remains bounded. For *ϵ > ρ*, one can construct contamination to force divergence, hence the breakdown point equals *ρ*.

#### Proposition B.5

(Scale Alignment via *κ*-Calibration). *The κ-calibration procedure re-scales the normalization factor of dGSEA to match that of classical GSEA. Specifically, the calibrated denominator becomes an estimator of the classical null expectation*, 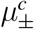. *This ensures that the resulting* dNES *values are on a scale directly comparable to classical* NES *values*.

*Proof*. Let 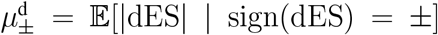 and 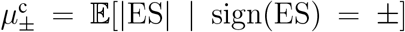 be the population-level robust means under the null. The calibration factor is defined as 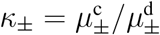. The calibrated denominator for dGSEA is 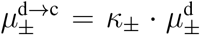. Substituting the definition of *κ*_*±*_ yields:

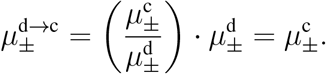

Therefore, the calibrated score 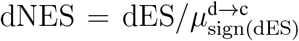 is normalized by an estimator of the same expected null value as classical GSEA. The two scores are thus expressed on the same statistical scale.

*Remark* B.2. As a direct consequence of normalization by the conditional mean, it holds that 𝔼[NES^classical^ | sign(ES) = ±] = 1 under the null. The *κ*-calibration ensures that dNES is normalized by the classical GSEA scale, making their values directly comparable.

### B.3 Differentiability and Gradient Properties

#### Proposition B.6

(Continuity, a.e. differentiability, and gradient bounds). *Assume* 1 ≤ |𝒮| ≤ *G* − 1 *and let* 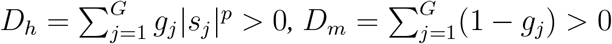. *If p* ≥ 1 *(or* |*s*|^p^ *is smoothed at s* = 0*), then* dES(*s, g*) *is continuous everywhere and differentiable almost everywhere in s. Moreover, writing σ*^*′*^(*z*) = *σ*(*z*)(1 − *σ*(*z*)) ≤ 1*/*4, *the following bounds hold for any i, ℓ:*

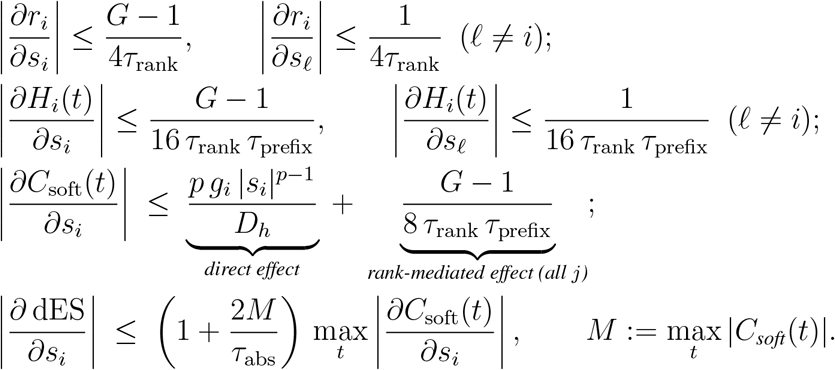

*Proof*. Continuity and almost-everywhere differentiability follow because all building blocks (*σ*, power function for *p* ≥ 1, summation) are continuous and smooth, except for the absolute value function | · | used in the softmax weights, which is non-differentiable only on a null set where *C*_soft_(*t*) = 0. The gradient bounds are derived by repeated application of the chain rule and the bound *σ*^*′*^ ≤ 1*/*4. The rank-mediated effect on *C*_soft_(*t*) arises from differentiating 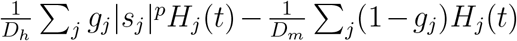 with respect to *s*_*i*_. Using the triangle inequality and the bounds |*∂H*_*j*_*/∂s*_*i*_| ≤ 1*/*(16*τ*_rank_*τ*_prefix_) for *j* ≠ *i* and |*∂H*_*i*_*/∂s*_*i*_| ≤ (*G* − 1)*/*(16*τ*_rank_*τ*_prefix_), the total contribution is bounded by (*G* − 1)*/*(8*τ*_rank_*τ*_prefix_). The final bound for *∂* dES*/∂s*_*i*_ results from differentiating the softmax-weighted sum, where the Jacobian of the softmax weights contributes the factor proportional to *M/τ*_abs_.

### B.4 Acceleration

#### Theorem B.7

(Nyström Approximation Error Bound – Robust Version). *Let F*_*G*_ *be the empirical distribution of* {*s*_1_, …, *s*_*G*_} *and F*_*A*_ *the empirical distribution of anchors* 𝒜_*m*_ = {*a*_1_, …, *a*_*m*_}.

*For a fixed i, set f*_*i*_(*x*) = *σ*((*x* − *s*_*i*_)*/τ*_rank_), *and denote the exact and approximated soft ranks by*

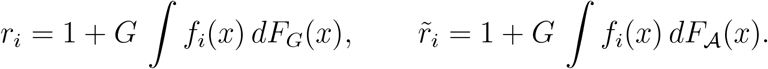

*Then*

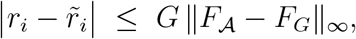

*where* ∥*F*_*A*_ −*F*_*G*_∥_∞_ := sup_*x*∈ℝ_ |*F*_𝒜_(*x*)−*F*_*G*_(*x*)| *is the (Kolmogorov–Smirnov) star-discrepancy w*.*r*.*t. F*_*G*_.

*Proof*. By definition 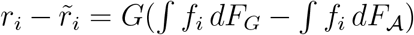. Since *f*_*i*_ is monotone increasing with total variation Var(*f*_*i*_) = *f*_*i*_(+∞) − *f*_*i*_(−∞) = 1, Stieltjes integration by parts gives

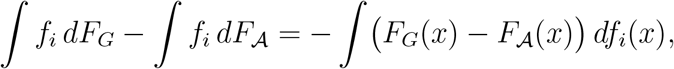

whence

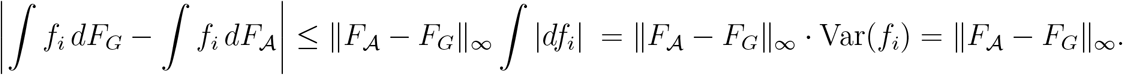

Multiplying by *G* gives the bound.

#### Proposition B.8

(Windowed Grid Error for dES; Exactness Conditions). *Let M* = max_*t*_ |*C*_*soft*_(*t*)| 𝒯 ⊂ {1, …, *G*} *be the windowed grid, and define the softmax weights on the full grid w*_*t*_ = exp( | *C*_*soft*_ (*t*) | */τ*_abs_ )*/* ∑ _u_ exp( | *C*_*soft*_ (*u*) | */τ*_abs_ ), *with outside-window mass ρ* := ∑ _t∉𝒯_ *w*_t_ ^∈^ [0, 1). *Let* dES_full_ = ∑ _t_ *w*_*t*_*C*_*soft*_(*t*) *and* dES_win_ = ∑ _t∈𝒯_ *w*_*t*_*/*(1 − *ρ*) *C*_*soft*_(*t*) *(the same weights renormalized on* 𝒯 *). Then*

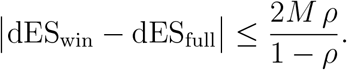

*Moreover, if the window contains* all *maximizers of* |*C*_*soft*_(*t*)| *and δ*_𝒯_ := min_*t*∉𝒯_ {*M* −|*C*_*soft*_(*t*)|} *>* 0, *then*

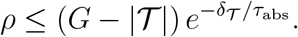

*Consequently, as τ*_abs_ ↓ 0 *(or δ*_𝒯_ ↑ ∞*), ρ* → 0 *and* dES_win_ → dES_full_.

*Proof*. Let *S*_in_ = ∑_t∈𝒯_ exp(|*C*_*t*_|*/τ*_abs_) and *S*_out_ = ∑_t∉𝒯_ exp(|*C*_*t*_|*/τ*_abs_), so *ρ* = *S*_out_*/*(*S*_in_ + *S*_out_). For *t* ∈ 𝒯, the re-normalized weight is *w*_*t*_*/*(1 − *ρ*). Then

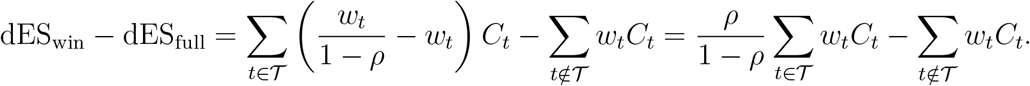

By the triangle inequality, 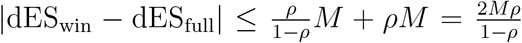. For the second claim, let *t*^⋆^ ∈ arg max_*t*_ |*C*_*t*_| be in 𝒯 . Then 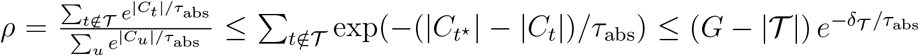.

#### Theorem B.9

(Time and Space Complexity). *Let B be the batch size, G the number of genes, m the number of Nyström anchors, T* = |𝒯 | *the size of the prefix grid (window), and K the number of permutations. With a batched tensor implementation and permutation chunk size c, the asymptotic costs are:*

`orig` *Soft rank via all-pairs: building H* ∈ ℝ^B*×*G*×*G^ *costs* Θ(*BG*^2^) *flops and stores* Θ(*BG*^2^) *elements; the smooth prefix step on the full grid is also O*(*BG*^2^).

`nystrom` *Soft rank: computing H* ∈ ℝ^B*×*G*×*m^ *costs* Θ(*BGm*) *flops with peak memory of O*(*BG* + *Bm*) *using streaming. The full-grid prefix step dominates time at O*(*BG*^2^).

`nyswin` *Combining Nyström rank and windowed prefix: per evaluation the cost is O*(*BGm*+ *BGT* ). *The full procedure with K permutations costs*

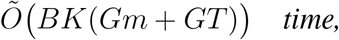

*and the peak memory is*

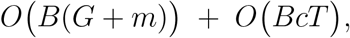

*where the second term is the working buffer for c permutations of the prefix curve*.

*Proof*. The analysis assumes that the cost of computing the *m* empirical quantile anchors, typically *O*(*G*) using efficient selection algorithms, is negligible compared to the dominant terms. The (`orig`) complexity follows from materializing the pairwise tensor. For (`nystrom`), *m*-anchor comparisons replace *G*-anchor comparisons, and memory can be optimized by streaming over anchors to avoid storing the full *B* × *G* × *m* tensor. For (`nyswin`), the windowed prefix step evaluates *H*_*i*_(*t*) for *t* ∈ 𝒯 only, costing *O*(*BGT* ). The time for the full procedure sums the per-permutation evaluation costs over *K* permutations. The peak memory is determined by storage for inputs and ranks (*O*(*B*(*G* + *m*))) plus the working buffer for one chunk of *c* permutations (*O*(*BcT* )).

## C Hyperparameter robustness and recommended defaults

To assess the practical usability of dGSEA in training-time settings, we examine the sensitivity of the enrichment scores to its three temperature parameters: *τ*_rank_ (soft ranking), *τ*_prefix_ (smooth prefix accumulation), and *τ*_abs_ (softmax aggregation over prefix positions). In contrast to downstream task performance, the focus here is on the stability of enrichment *semantics*— specifically, agreement with classical GSEA—under moderate parameter variation.

### C.1 Grid sweep and robustness analysis

We perform a grid sweep over the following ranges: *τ*_rank_ ∈ [0.8, 1.6], *τ*_prefix_ ∈ [0.8, 1.2], and *τ*_abs_ ∈ {0.40, 0.60, 0.80, 1.00}. For each configuration, we compute the Spearman correlation between normalized enrichment scores (NES) from classical GSEA and the corresponding dNES from dGSEA across representative synthetic scenarios. As shown in Fig. 10, dNES– NES correlations remain above 0.9 over a broad region of the parameter space, indicating a wide plateau of high semantic agreement and limited sensitivity to precise hyperparameter tuning.

**Figure 10:**
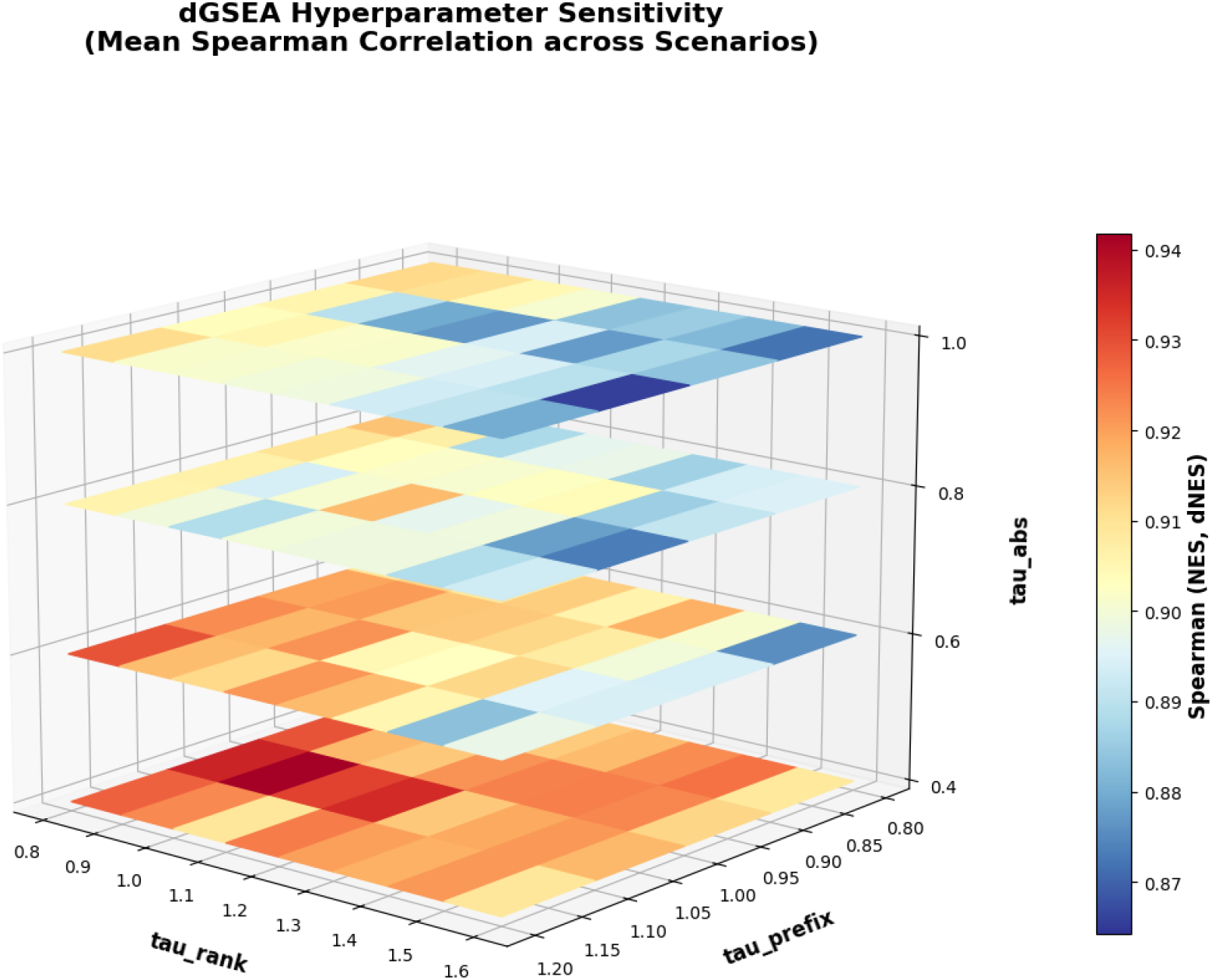
Hyperparameter sensitivity of dGSEA. The strongest effect is observed for *τ*_abs_, while a broad high-correlation plateau is present across *τ*_rank_ and *τ*_prefix_.

### C.2 Interpretation of parameter roles

The sweep clarifies the functional roles of the three parameters. The ranking temperature *τ*_rank_ primarily controls the sharpness of the soft ranking operation and exhibits a relatively flat correlation ridge over a wide range, indicating low sensitivity. The prefix temperature *τ*_prefix_ has minimal impact, reflecting the inherent robustness of the running-sum accumulation to moderate smoothing. In contrast, the aggregation temperature *τ*_abs_ exerts the strongest influence by modulating the softmax weighting over prefix positions: excessively small values over-emphasize local extrema, whereas intermediate values balance sensitivity and numerical stability.

### C.3 Recommended defaults

Aggregating results across scenarios yields a representative operating point at (*τ*_rank_, *τ*_prefix_, *τ*_abs_) (0.80, 1.10, 0.70) (Table 8), with contiguous “safe ranges” around this setting in which the mean Spearman correlation remains within ≈ 0.01 of the best observed value. These defaults are used throughout the experiments unless otherwise stated. We emphasize that this choice is not tuned to maximize any downstream task performance, but rather reflects a stable configuration that preserves enrichment semantics while avoiding excessive sensitivity to local extrema.

**Table 7:** Directional pathways used on LINCS L1000.

| Pathway | Representative process | $n_{\text{up}}$ | $n_{\text{down}}$ | Missing genes |
| --- | --- | --- | --- | --- |
| P53_pathway | p53 tumor-suppressor response | 6 | 1 | None |
| CellCycle_CDK | Cell-cycle core (CDK/Cyclins) | 1 | 4 | None |
| HeatShock_DNAJ | Heat-shock response (DNAJ/HSP40) | 4 | 0 | DNAJB2 |
| Ubiquitin_DUB | Ubiquitin-proteasome (DUBs) | 4 | 0 | None |
| DNA_damage_GADD45 | DNA-damage (GADD45 family) | 2 | 0 | None |

**Table 8:** Aggregated recommendation from the hyperparameter sweep (mean over scenarios).

| Rank | $\tau_{\text{rank}}$ | $\tau_{\text{prefix}}$ | $\tau_{\text{abs}}$ | Mean Spearman (NES, dNES) |
| --- | --- | --- | --- | --- |
| 1 (Representative setting) | 0.80 | 1.10 | 0.70 | 0.919 |
| *Safe ranges (global mean within $\approx 0.01$ of best): $\tau_{\text{rank}} \in [0.80, 1.10]$, $\tau_{\text{prefix}} \in [0.80, 1.20]$.* |  |  |  |  |

### C.4 Annealing trajectories

For completeness, Fig. 11 illustrates representative annealing trajectories for the temperature parameters under multiple baselines. These trajectories are provided as implementation guidance and are not required for the semantic validity or stability results reported in the main text.

**Figure 11:**
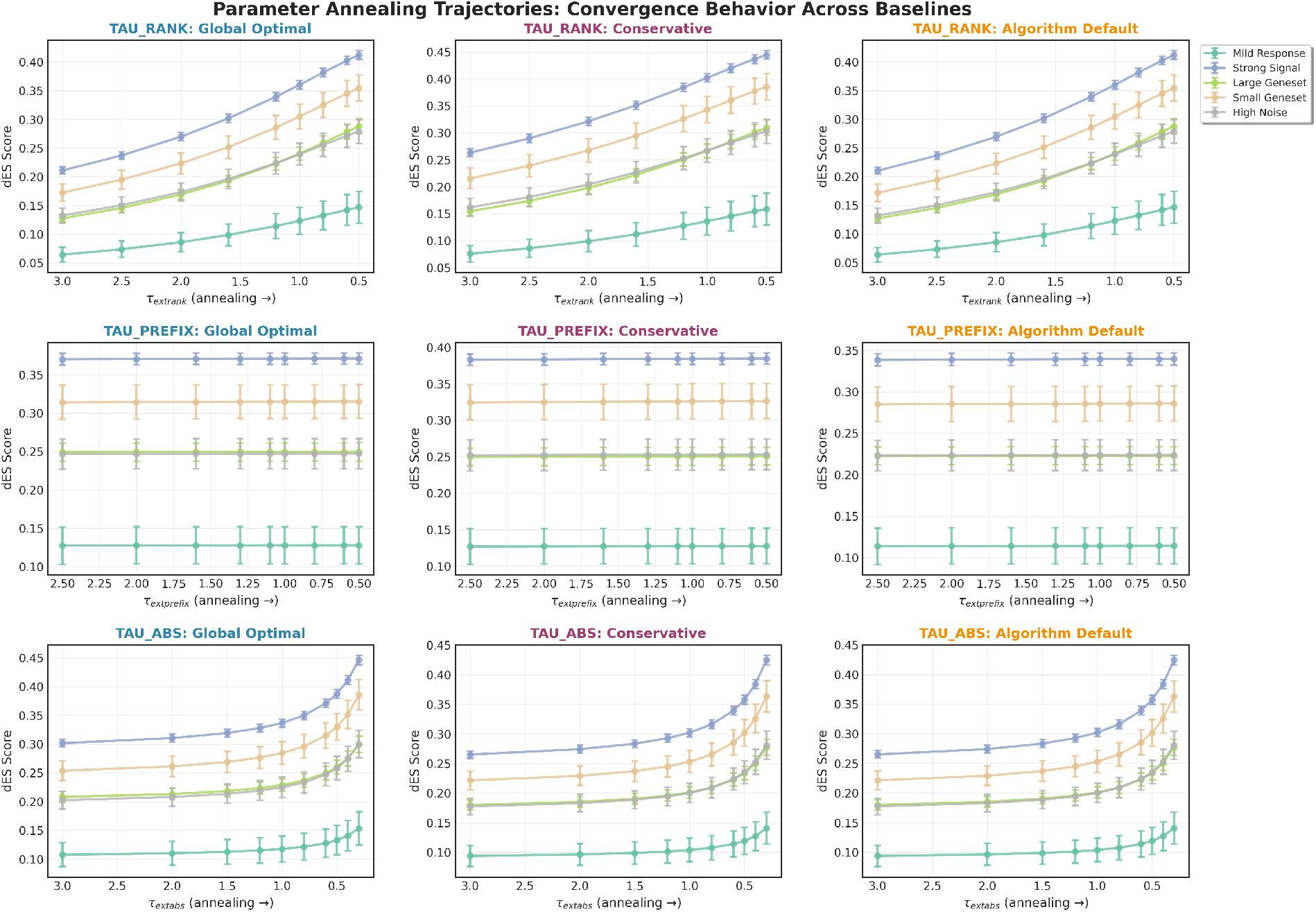
Representative temperature annealing trajectories used in selected experiments.

## Notes

### Competing Interest Statement

The authors have declared no competing interest.

